# Inhibition of CGRP receptor ameliorates AD pathology by reprogramming lipid metabolism through HDAC11/LXRβ/ABCA1 signaling

**DOI:** 10.1101/2025.10.17.683079

**Authors:** Guangchun Fan, Fangzheng Chen, Min Liang, Ping Yang, Hongtian Dong, Jing Su, Mengqi Wang, Ziyuan Wang, Penggang Ning, Chenye Shen, Xiwen Tang, Xin Yan, Haoxiang Lin, Jiayin Zhao, Yunhe Zhang, Zhesu Cheng, Hui Lu, Xiuping Liu, Renyuan Zhou, Yang Luan, Li Cao, Mei Yu, Wensheng Li, Jian Fei, Ruling Shen, Fang Huang

**Author notes:** Correspondence to: Dr. Mei Yu, Institutes of Brain Science, State Key Laboratory of Medical Neurobiology, Fudan University.; Dr. Wensheng Li, Department of Anatomy, Histology and Embryology, School of Basic Medical Sciences, Fudan University. E- mail; Dr. Jian Fei, School of Life Sciences and Technology, Tongji University.; Dr. Ruling Shen, Shanghai Laboratory Animal Research Center., or Dr. Fang Huang, Institutes of Brain Science, State Key Laboratory of Medical Neurobiology, Fudan University. These authors contribute equally (Guangchun Fan, Fangzheng Chen, Min Liang, Ping Yang, Hongtian Dong).

## Abstract

The Calcitonin gene-related peptide (CGRP) receptor has gained attention in Alzheimer’s Disease (AD) research due to its involvement in regulating neuroinflammation. However, its role and mechanism in AD pathology remain unclear. Here, we demonstrate that CALCRL, a core component of the CGRP receptor, is upregulated in the hippocampus of AD dementia patients and 5×FAD mice. Knockout of the CGRP receptor ligand Calca or pharmacological blockade using Rimegepant (Rim), reduces soluble Aβ1-42 oligomer-induced neuronal death and glial inflammation. Rim treatment also rescues neurobehavioral impairments, neurodegeneration, and lipid metabolism dysfunction in 5×FAD mice. Mechanistically, these effects are mediated through HDAC11 inhibition, which enhances LXRβ acetylation and ABCA1 expression, promoting the reprogramming of neuronal lipid metabolism. Importantly, this CALCRL/HDAC11/LXRβ/ABCA1 axis is conserved across both humans and mice. Our findings uncover a novel mechanism underlying AD pathogenesis and highlight the therapeutic potential of targeting CGRP signaling in AD.

**HIGHLIGHTS:** Inhibition of CGRP receptor ameliorates disease pathology in models of AD HDAC11 mediates CGRP receptor function in AD

HDAC11 is a pivotal regulator in lipid metabolism by LXRβ/ABCA1 signaling

The HDAC11/LXRβ/ABCA1 axis is conserved in AD humans and mice

## INTRODUCTION

Alzheimer’s disease (AD) is the leading cause of dementia worldwide, characterized by multiple pathological features in the brain, including amyloid-β (Aβ) plaques, tau tangles, neuroinflammation, and synaptic terminal damage (Jucker and Walker, 2023; Masters et al., 2015; Scheltens et al., 2021). However, strategies to slow or prevent its clinical progression have remained elusive until recently. Therefore, identifying new and effective therapeutic targets is critical.

The Calcitonin Gene-Related Peptide (CGRP) receptor is a complex composed of multiple proteins, with two critical transmembrane proteins—calcitonin receptor-like receptor (CALCRL) and receptor activity-modifying protein 1 (RAMP1)—playing key roles in mediating the effects of CGRP (Brain and Grant, 2004; Russell et al., 2014). CGRP is a 37-amino acid neuropeptide that exists in two forms, α and β CGRP, with the α CGRP predominantly transcribed from the calcitonin– CGRP gene (CALCA) and playing a major role in the central nervous system (CNS) (Edvinsson et al., 2018). CGRP and its receptors are present in several regions of the CNS, including the dorsal horn of the spinal cord, the trigeminal nucleus caudalis, the dorsal and trigeminal ganglia, the periaqueductal gray, the amygdala, and the thalamus (Hay and Walker, 2017; Iyengar et al., 2017). The distribution of CGRP and its receptor in these regions highlights their pivotal involvement in pain modulation, neuroinflammatory processes, and neurovascular regulation, particularly in conditions such as migraine and other neurovascular disorders (Edvinsson, 2019; Ho et al., 2010). While neuroinflammation represents a key pathological driver in both the initiation and progression of AD, the specific role and underlying mechanism of the CGRP receptor signaling in AD remain to be elucidated.

In the present study, we compared the expression levels of CALCRL between AD patients and normal brain tissue across different single-cell databases and found that CALCRL was upregulated in AD samples. These findings were confirmed both in hippocampal tissue from AD dementia patients and 5×FAD mice, a widely used transgenic model of AD. Next, we examined whether inhibition of CGRP receptor function could influence AD pathology. *In vitro*, inhibition of CGRP receptor activity through ligand knockout (*Calca^-/-^*) or pharmacological blockade using the receptor antagonist Rimegepant (Rim, a clinically approved drug for migraine treatment) attenuated sAβ1- 42 oligomer-induced neuronal death and suppressed glial-mediated inflammatory responses. *In vivo*, 5×FAD mice at 7 months of age that received intraperitoneal injections of Rim every other day for one month exhibited significant improvements in anxiety-like behaviors and spatial memory deficits. This treatment also mitigated several AD-related pathologies, including amyloid plaque deposition, tau hyperphosphorylation, neuronal death, neuroinflammation, and lipid metabolism dysfunction, particularly in the hippocampus, without significant sex differences. To further explore the underlying mechanisms, we integrated multi-omics data from the hippocampus of mice, and demonstrated that the therapeutic efficacy of Rim in AD is attributable to the inhibition of HDAC11. Notably, knockdown of *Hdac11* for up to one month in 5×FAD hippocampal neurons recapitulated the beneficial effects of Rim treatment. Finally, we demonstrated that the protective effect of HDAC11 inhibition in AD depends on the maintenance of LXRβ acetylation, which enhances ABCA1 expression. This molecular cascade is further supported by the observation that HDAC11 is strongly upregulated in AD dementia patients and 5×FAD mice, and its expression is negatively correlated with ABCA1 levels. Our study uncovered a novel role of CGRP receptor in regulating lipid metabolism and neuroinflammation via the HDAC11/LXRβ/ABCA1 axis, highlighting a potential therapeutic strategy for AD.

## RESULTS

### CALCRL is upregulated in patients with Alzheimer’s dementia and 5×FAD mice

To investigate the expression pattern of CALCRL in AD, we queried the Seattle Alzheimer’s Disease Brain Cell Atlas (SEA-AD) database, which includes single-nucleus RNA sequencing (snRNA-Seq) data from the middle temporal gyrus (MTG) of 33 male and 51 female donors (Figure 1A) (Gabitto et al., 2024). The results revealed that CALCRL was highly expressed in multiple cell types in both the older dementia and no-dementia groups, compared to reference genomes from younger individuals (Figure 1B-D and S1A) (Jorstad et al., 2023). Additionally, we retrieved occipital cortex snRNA-Seq data from the NCBI database (https://www.ncbi.nlm.nih.gov/) using the accession code GSE148822, which includes data from 3 male and 7 female AD donors (Figure S1B) (Gerrits et al., 2021). Consistent with the SEA-AD database results, CALCRL expression was similarly upregulated in older samples compared with reference genomes from younger individuals (Figure S1C- S1G). To further evaluate CALCRL protein expression, we conducted Western blot analysis on hippocampal samples from both male (n=3) and female (n=3) dementia and non-dementia participants (Figure 1E and S1H). The results show that CALCRL protein level was significantly increased in dementia patients compared to no-dementia controls (Figure 1F and 1G). Similarly, we examined CALCRL expression in the hippocampus of 5×FAD mice and obtained consistent findings: CALCRL was upregulated in both female and male 5×FAD mice compared to wild-type (WT) controls (Figure 1F and 1H). These results suggest that the expression and function of CALCRL may play an important role in the pathological processes of AD.

**Figure 1.**
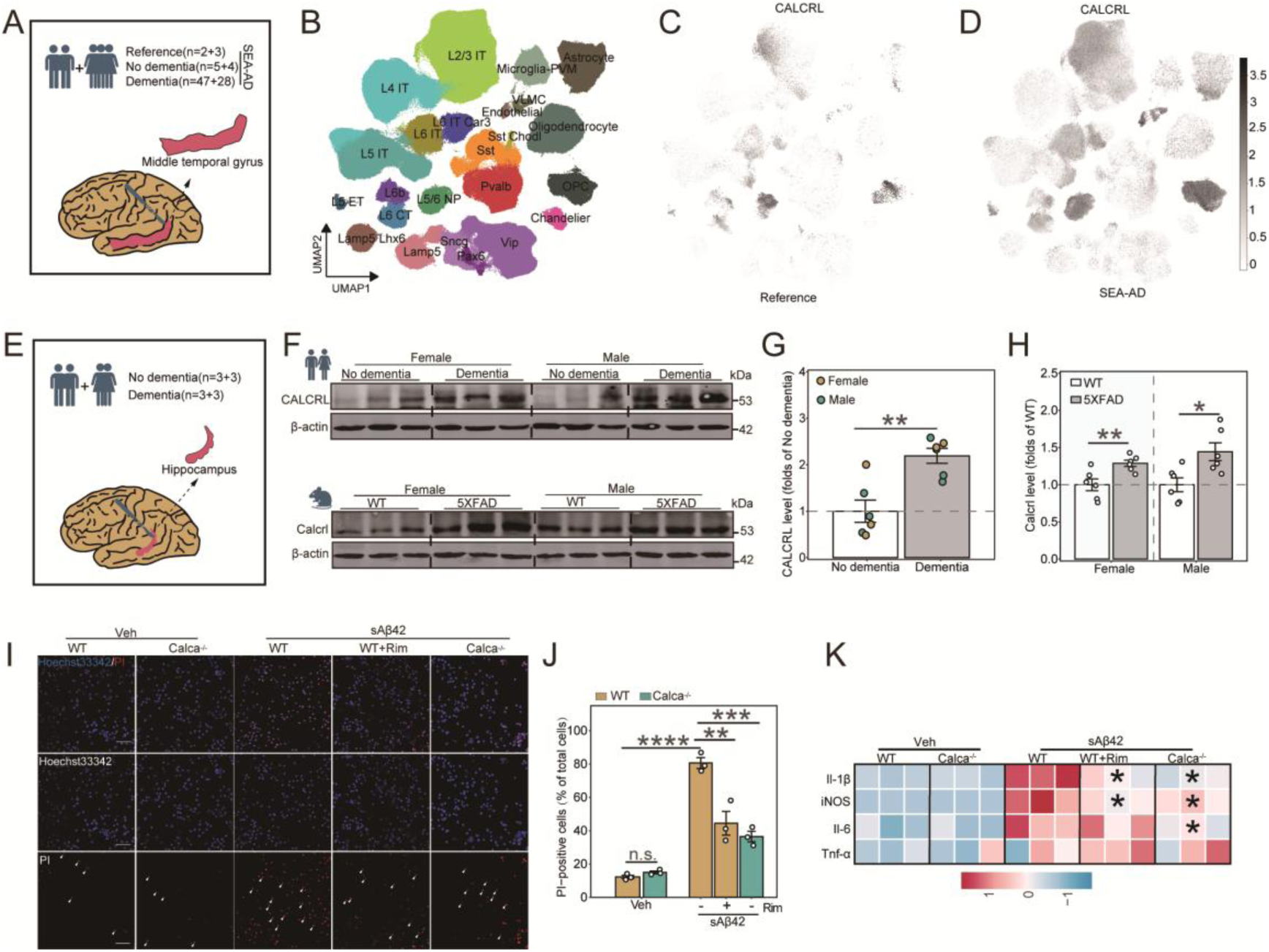
Inhibition of the CGRP receptor reduces primary neuronal cell death and glial inflammation induced by sAβ1-42. (A) Characteristic information and the number of subjects in each group. (B) UMAP of the associated cell type and clusters. VLMC, vascular/leptomeningeal cells; OPC, oligodendrocyte precursor cells; L5/6 NP, layer 5/6 near-projecting neurons; L6 IT Car3, layer 6 intratelencephalic neurons (Car3⁺); L6b, layer 6b excitatory neurons; L6 CT, layer 6 corticothalamic neurons; L5 ET, layer 5 extratelencephalic neurons; L5 IT, layer 5 intratelencephalic neurons; L4 IT, layer 4 intratelencephalic neurons; L6 IT, layer 6 intratelencephalic neurons; L2/3 IT, layer 2/3 intratelencephalic neurons; Sst, somatostatin⁺ interneurons; Vip, vasoactive intestinal peptide⁺ interneurons. (C and D) UMAP visualization of cells expressing CALCRL in the reference (C) and SEA-AD (D) groups. (E) Characteristic information and the number of subjects in each group. (F-H) Western blot image (F) and densitometry analysis of CALCRL in human (G) and mouse (H) hippocampus. (I and J) Representative images of Hoechst 33342 and PI double staining (I) and the quantification of PI-positive cells (J) in primary neurons from WT and *Calca* KO mice treated with Veh, sAβ1-42 or Rim + sAβ1-42. (K) Heatmap shows the expression of *Il-1β*, *iNOS*, *Il-6* and *Tnf-α* mRNA in primary mixglia stimulated by sAβ1-42. n=3-6/genotype/treatment. Scale bars: 50 μm in (I). \**p* < 0.05, \*\**p* < 0.01, \*\*\**p* < 0.001, \*\*\*\**p* < 0.0001. n.s., non-significant. Statistical analyses were analyzed by unpaired Student’s t test. See also Figure S1.

### Inhibition of CGRP receptor reduces primary neuronal cell death and glial inflammation induced by soluble Aβ1-42 oligomers

Initially, we conducted molecular docking simulations of Rim, a well-recognized CGRP-receptor antagonist, on the crystal structure of the ectodomain complex of the CGRP receptor (PDB: 3N7R).

The results revealed that Rim establishes two hydrogen bonds with THR122 and ARG119, respectively (Figure S1I). Accumulating evidence indicates that soluble Aβ1-42 oligomers (sAβ1- 42) are closely linked to the pathogenesis of AD. To investigate whether the CGRP receptor modulates neuronal and glial responses to sAβ1-42-induced injury, we prepared sAβ1-42 from synthetic mouse Aβ1-42 monomers, as described (Mhillaj et al., 2018). Scanning electron microscopy (SEM) images of purified fibrin clots revealed the formation of aggregates in the presence of Aβ1-42, suggesting that sAβ1-42 was successfully prepared (Figure S1K). Subsequently, we isolated and cultured primary cortical neurons and mixed glial cells from both WT and *Calca* knockout mice, and treated these cells with 10 μM sAβ1-42, with or without Rim, for 24 hours *in vitro* (Figure S1J and S1M). The viability of primary neurons deficient in *Calca* exhibited no significant difference when compared to that of WT neurons. In contrast, treatment with Rim at concentrations of 0.1 and 1 μM, as well as *Calca* knockout, significantly enhanced neuronal viability compared to the control group following sAβ1-42 exposure, as determined by the cell counting kit- 8 (CCK-8) assay. This protective effect was further corroborated by Hoechst 33342/propidium iodide (PI) staining (Figure 1I and 1J). Furthermore, the mRNA levels of proinflammatory factors, including IL-1β, iNOS, IL-6, and TNF-α, were measured in primary mixed glial cells treated with sAβ1-42 for 24 hours, with or without 1 μM Rim. Notably, a significant reduction in *Il-1β* and *NOS2* expression was detected in the Rim + sAβ1-42 group compared to the sAβ1-42 group. Additionally, in the *Calca*^-/-^ + sAβ1-42 group, the expression levels of *Il-1β*, *NOS2*, and *Il-6* were significantly downregulated compared to the WT + sAβ1-42 group (Figure 1K). Collectively, these findings highlight the potential of CGRP receptor inhibition as a promising therapeutic approach for the treatment of AD.

### CGRP receptor antagonist Rim improves behavioral performance in AD worms and 7-month-old 5×FAD mice

To further investigate the therapeutic potential of Rim in mitigating Aβ-induced pathology, we utilized both the transgenic *Caenorhabditis elegans* AD model (Punc-54::Aβ1–42, strain CL2006), which exhibits progressive adult-onset paralysis (Figure S1N), and the 5×FAD transgenic mouse model. As shown in Figure S1O, treatment with 2 μM Rim significantly delayed the onset of Aβ- induced paralysis in CL2006 nematodes. 5×FAD transgenic mice begin to develop Aβ deposition in the brain as early as 2 months of age (Eimer and Vassar, 2013). In this study, we used 7-month-old 5×FAD mice of both sexes, as these mice exhibit substantial accumulation of Aβ plaques. Since Rim orally disintegrating tablets (75 mg) are typically administered daily for acute migraine treatment and every other day for migraine prevention (Mullin et al., 2020), we adopted a regimen of alternate-day intraperitoneal injections of Rim (10mg/kg) in 5×FAD mice for a duration of one month (Figure 2A).

**Figure 2.**
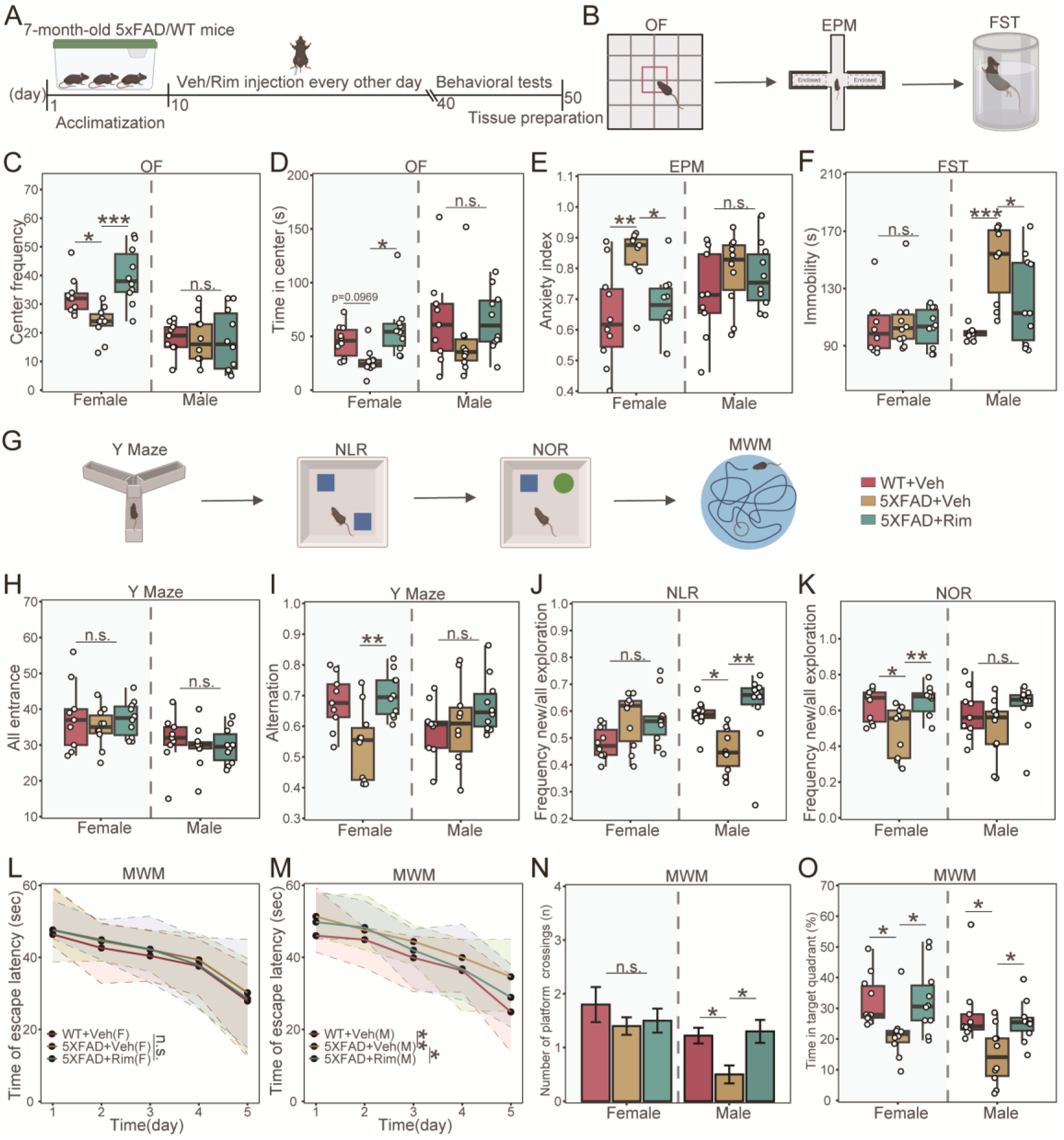
Rim improved behavioral deficits observed in AD mice (A) Schematic representation of the experimental design. (B) Cartoon schematic of the behavior experimental design. (C and D) The evaluated parameters from the Open field test (OF) included the number of times entering the central area and the residence time of the central area. (E) The anxiety index calculated by the Elevated plus maze (EPM). (F) Immobility time in the Forced swim test (FST). (G) Schematic of the behavior experimental design. (H and I) The number of total arm entries and spontaneous alternations in the Y-maze test. (J) The ratio of the number of explorations of the new location object to the total number of explorations in the Novel location recognition (NLR) task. (K) The ratio of the number of explorations of the new object to the total number of explorations in the Novel object recognition (NOR) task. (L-O) The results of Morris water maze (MWM). (L, M) Latency to the platform in female (L) and male (M); (N)Entries into the platform zone; (O) The percentage of time spent in target quadrant. n=9-10/genotype. \**p* < 0.05, \*\**p* < 0.01, \*\*\**p* < 0.001, n.s., non-significant. Statistical analyses were analyzed by one-way ANOVA followed by a post hoc Dunnett’s test.

Previous studies have reported that 5×FAD mice typically display pronounced anxiety- and depression-like behaviors between 4 to 6 months of age (Forner et al., 2021). In this study, we sought to evaluate the effects of Rim on such behavioral deficits in 5×FAD mice using the Open field test (OF), Elevated plus maze (EPM), and Forced swim test (FST) (Figure 2B). Our findings revealed that Rim treatment significantly increased both the frequency and duration of entries into the central area in the OF (Figure 2C and 2D), indicating reduced anxiety-like behavior. Additionally, Rim significantly lowered the anxiety index in the EPM, particularly in female 5×FAD mice (Figure 2E). Furthermore, in male 5×FAD mice, Rim treatment markedly reduced immobility time in the FST (Figure 2F), suggesting an antidepressant-like effect.

The effects of Rim at the selected doses on learning and memory were further evaluated using a series of behavioral tests (Figure 2G). The Y-maze test, which is partially linked to the septo- hippocampal system involved in learning and spatial working memory (Kim et al., 2023), revealed that Rim treatment significantly and substantially reversed the alternation deficits observed in female 5×FAD mice (Figure 2H and 2I). However, in male 5×FAD mice, the improvement following Rim treatment did not reach statistical significance (Figure 2H and 2I). To further assess the impact of Rim on memory, we employed the Novel location recognition (NLR) and Novel object recognition (NOR) tests, two tasks influenced by hippocampus- and cortex-mediated alterations (Drulis-Fajdasz et al., 2023) (Figure 2G). Male 5×FAD mice exhibited deficits in NLR, while female 5×FAD mice displayed impairments in NOR (Figure 2H and 2I). Notably, Rim treatment significantly restored the ability of male mice to recognize novel locations and female mice to recognize novel objects (Figure 2H and 2I). Finally, the Morris water maze (MWM) test was used to examine hippocampus-dependent spatial memory, a cognitive domain known to be affected early in AD (Curdt et al., 2022) (Figure 2G). During the training phase, Rim-treated male 5×FAD mice demonstrated a significantly reduced latency to find the platform compared to Veh-treated mice on day 5 (Figures 2L and 2M). In the test phase, Rim-treated male mice also exhibited more platform crossings and less time spent outside the platform quadrant compared to Veh-treated animals (Figures 2N and 2O). In contrast, Rim-treated female 5×FAD mice displayed a significant increase in the time spent in the platform quadrant during the test phase, but no differences were observed in other measures (Figure 2N and 2O). These findings suggest that Rim exerts sex-dependent effects on learning and memory, with notable improvements in specific cognitive domains across both male and female 5×FAD mice.

### CGRP receptor antagonist Rim alleviates AD-related pathology

Rim demonstrated its potential to alleviate cognitive deficits in 5×FAD mice of both sexes. To explore the mechanisms underlying Rim’s beneficial effects on cognitive function, we investigated several key pathological features associated with AD, including the amyloid-β (Aβ) production, tau pathology, and synaptic abnormalities in the brains of 5×FAD mice. Using Thioflavin-S staining, we quantified the number and size of Aβ plaques in the cortex and hippocampus respectively. As expected, 5×FAD mice exhibited a significant increase in Aβ plaques compared to WT controls (Figure 3A-C and S2A-C). Notably, Rim administration led to a sex-specific reduction in Aβ pathology. In female 5×FAD mice, Rim treatment resulted in more than a 50% decrease in plaque number and total plaque area in the hippocampus (Figure 3B and 3C), along with a significant reduction in total plaque area in the cortex (Figure S2C). In male 5×FAD mice, Rim treatment significantly reduced the total plaque area in the hippocampus, though the effect was less pronounced compared to females (Figure 3C). Overall, these results suggest that Rim mitigates Aβ pathology in a sex-dependent manner, with more robust effects observed in female 5×FAD mice. This reduction in Aβ burden may contribute to the cognitive improvements observed in Rim-treated animals.

**Figure 3:**
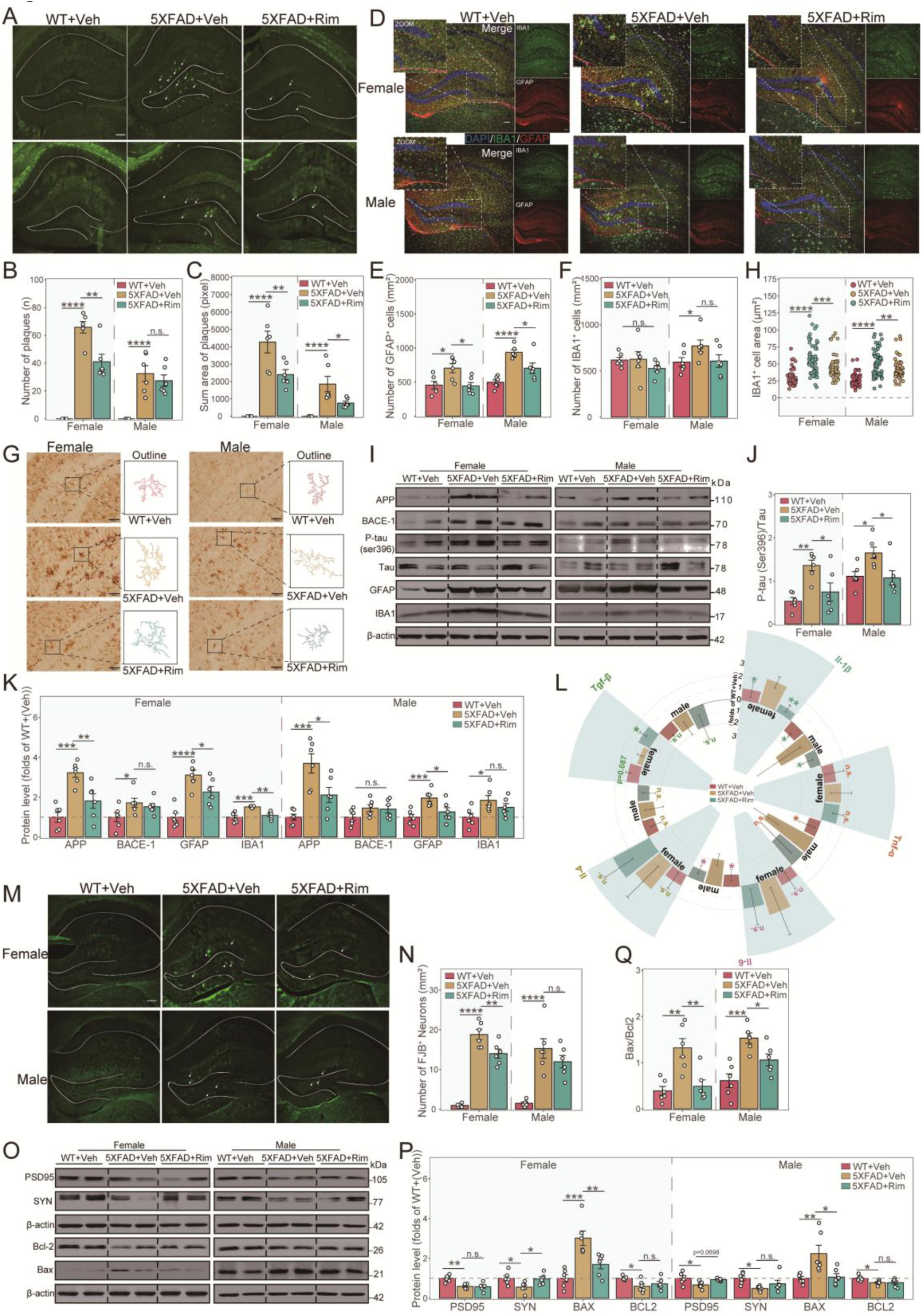
Rim alleviates AD-related pathological features in the hippocampus (A) Representative Thioflavin-S staining hippocampal images of WT and 5×FAD mice after the administration of the vehicle or Rim. (B and C) Quantitative measurements of Aβ plaque numbers (B) and area (C) in the hippocampus. (D) Immunofluorescence staining of GFAP (red) and IBA-1 (green) in the hippocampus. (E and F) Quantifications of GFAP^+^ astrocytes (E) and IBA-1^+^ microglia (F) in the hippocampus. (G and H) Representative images of IBA-1 immunostaining (G) and the quantified IBA-1^+^ cell body volume (H) in the hippocampus. (I-K) Western blot image (I) and densitometry analysis of AD-related marker proteins including APP, BACE1, phosphorylated tau, total tau, GFAP and IBA-1 in the hippocampus (J and K). (L) Quantitative PCR analysis of pro-inflammatory factors *Il-1β*, *Tnf-α*, and *Il-6*, and anti-inflammatory factors *Il-4* and *Tgf-β* expression in the hippocampus. (M) Representative Fluoro-Jade B staining images. (N) Quantitative measurements of FJB^+^ neuron numbers. (O-Q) Western blot image (O) and densitometry analysis of PSD95, synaptophysin (SYN), Bcl-2 and Bax in the hippocampus (P and Q). n=3-6/genotype; n=60 IBA-1^+^ cell, 6/group (H). Scale bar: 100 μm in (A, B and M); 200 μm in (G). \**p* < 0.05, \*\**p* < 0.01, \*\*\**p* < 0.001, \*\*\*\**p* < 0.0001, n.s., non-significant. Statistical analyses were analyzed by one-way ANOVA followed by a post hoc Dunnett’s test. See also Figure S2.

The above findings were further supported by the effects of Rim on amyloid precursor protein (APP) levels. Rim-treated 5×FAD mice exhibited significantly reduced APP levels in the hippocampus, while only a decreasing trend was observed in the cortex (*p* = 0.0781) (Figure 3I and 3K). Given that Aβ formation is a hallmark of AD, we further examined the expression of β-site amyloid precursor protein cleaving enzyme 1 (BACE1), a key enzyme in APP processing (Hur, 2022). BACE1 protein levels were significantly elevated in 5×FAD mice compared to WT controls (Figure 3I, 3K, S2I and S2K). However, no significant reduction in BACE1 levels was detected between Rim- and vehicle (Veh)-treated 5×FAD mice (Figure 3I, 3K, S2I and S2K). Hyperphosphorylated tau (p-tau) is another critical pathological feature of AD, as tau hyperphosphorylation triggers pathological changes, including synaptic impairment (Khurana et al., 2006). We thus assessed p-tau burden to evaluate Rim’s effects on tau pathology. By analyzing the expression level of p-tau (S396), we observed a significant reduction in p-tau levels in the hippocampus of Rim-treated 5×FAD mice (Figure 3I and 3J). To further investigate synaptic alterations, we examined the pre- and post- synaptic proteins associated with synaptic plasticity and memory deficits (Xu et al., 2021). The expression levels of post-synaptic density protein 95 (PSD95) and pre-synaptic protein synaptophysin (SYN) remained unchanged in the cortex of 5×FAD mice of both sexes (Figure S2O and S2P). However, PSD95 proteins exhibited an increasing trend in the hippocampus of Rim- treated male 5×FAD mice (Figure 3O and 3P); In contrast, Rim treatment significantly upregulated synaptophysin levels in the hippocampus of female 5×FAD mice (Figure 3O and 3P). Our findings demonstrate that Rim treatment alleviates Aβ accumulation and synaptic dysfunction, and also inhibits the hyperphosphorylation of tau proteins, particularly in the hippocampus. These results highlight Rim’s potential therapeutic effects on multiple pathological features of AD.

The proliferation and activation of microglia, particularly those surrounding amyloid plaques, are remarkable characteristics of AD (Leng and Edison, 2021). Activated microglia can aggravate tau pathology and release inflammatory factors that directly harm neurons or do so indirectly through the activation of neurotoxic astrocytes (Kwon and Koh, 2020). To assess the impact of Rim on neuroinflammation in AD brains, we evaluated changes in astrocytosis and microgliosis through immunofluorescence staining and Western blots analysis. Specifically, we examined the expression of glial fibrillary acidic protein (GFAP) and ionized calcium-binding adaptor molecule 1 (IBA-1), respectively. Our results, as depicted in Figure 3D-F and S2D-F, revealed an increase of microglia and astrocytes in the hippocampus and cortex of 5×FAD mice compared to their WT counterparts, with a noted decrease in astrocytes following Rim administration. Furthermore, through IBA-1 immunohistochemistry staining and morphological analysis, we observed that microglia surrounding plaques in 5×FAD mice exhibited an amoeboid shape with somatic hypertrophy, whereas Rim-treated microglia generally displayed reduced cell bodies (Figure 3G, 3H, S2G and S2H). Corroborating the findings of immunofluorescence staining, Rim treatment significantly downregulated GFAP and IBA-1 protein levels both in the cortex and hippocampus of female 5×FAD mice (Figure 3I, 3K, S2I and S3K). However, in male 5×FAD mice, Rim treatment only led to a decrease in GFAP expression in the hippocampus, with IBA-1 showing a downward trend without reaching statistical significance (Figure 3I, 3K, S2I and S2K).

Additionally, we evaluated the RNA expression levels of inflammation-related cytokines in the hippocampus and cortex of 7-month-old 5×FAD mice. Rim treatment significantly reduced *Il-1β* level in the hippocampus of both female and male 5×FAD mice and increased *Tgf-β* level in female 5×FAD mice, while *Il-6* level significantly decreased in male 5×FAD mice (Figure 3L). In the cortex, Rim treatment significantly reduced *Il-6* expression and increased *Tgf-β* level in female 5×FAD mice (Figure S2L), whereas no significant changes were observed in male 5×FAD mice (Figure 3L and S2L). Altogether, these findings suggest that Rim treatment mitigates neuroinflammation and modifies the microglial and astrocytic proliferation in 7-month-old 5×FAD mice, highlighting its potential therapeutic efficacy in suppressing neuroinflammation in AD.

Fluoro-Jade B, a widely recognized marker for identifying damaged and dying neurons, was employed to evaluate the neuroprotective effects of Rim in the hippocampus and cortex of 5×FAD mice (Ikenari et al., 2021). Our findings revealed a significant increase in the number of degenerated neurons within the cortex of 5×FAD mice when compared to the WT controls (Figure 3M, 3N, S2M and S2N). Notably, treatment with Rim resulted in a marked reduction in the FJB-positive cell numbers both in the hippocampus and cortex of female 5×FAD mice compared to those receiving vehicle treatment. In contrast, male 5×FAD mice exhibited only a decreasing trend in the hippocampal degeneration, which did not reach statistical significance (Figure 3M, 3N, S2M and S2N).

The pro-apoptotic protein Bax and the anti-apoptotic protein Bcl-2 are pivotal regulators of the mitochondrial pathway of apoptosis (Czabotar and Garcia-Saez, 2023). The impact of Rim on the expression of Bcl-2 and Bax in the brains of 5×FAD mice was further explored by Western blot analysis. The results indicated that in comparison to the WT control group, vehicle-treated 5×FAD mice exhibited a significantly reduced expression of Bcl-2, an elevated expression of Bax, and an increased Bax/Bcl-2 ratio in the hippocampus (Figure 3O-3Q). In the cortex, Bax expression levels and the Bax/Bcl-2 ratio showed a significant increase, while Bcl-2 expression remained unaltered (Figure S2O-Q). Notably, Rim treatment significantly reversed the changes in Bax levels and reduced the Bax/Bcl-2 ratio in the hippocampus (Figures 3O-3Q), but not in the cortex (Figure S2O- S2Q), demonstrating the neuroprotective effects of Rim specifically in the hippocampus of 7-month-old 5×FAD mice.

### Multi-omics analysis shows HDAC11 downregulation is key to the protective effects of CGRP receptor antagonist Rim in the hippocampus of 7-month-old 5×FAD mice

To delve deeper into the mechanisms underlying the impact of Rim on AD pathology, we conducted a comprehensive multi-omics analysis. Initially, RNA sequencing (RNA-seq) analysis was carried out on hippocampal tissues from 5×FAD mice, as depicted in Figure S3A. Applying an adjusted *p*- value threshold of < 0.05, we identified a set of differentially expressed genes (DEGs): 955 DEGs between 5×FAD+Veh(F) and WT+Veh(F) (Figure S3B), 717 DEGs between 5×FAD+Veh(M) and WT+Veh(M) (Figure S3C), 1053 DEGs between 5×FAD+Rim(F) and WT+Veh(F) (Figure S3D), and 591 DEGs between 5×FAD+Rim(M) and WT+Veh(M) (Figure S3E). Functional annotation of the DEGs, considering both Rim treatment and AD condition, was performed using Gene Ontology (GO) analysis. The results revealed that the most significantly impacted biological processes were related to immune responses (Figure S3F-S3I). Gene Set Enrichment Analysis (GSEA) was employed to identify subtle yet significant expression changes across gene sets linked by common pathway functions, which might not be discernible through direct gene comparison. Through GSEA, we examined inflammatory pathways as cataloged in the Kyoto Encyclopedia of Genes and Genomes (KEGG). As shown in Figure S3J, the analysis highlighted significant enrichment of several KEGG inflammatory pathways across the four groups, including the TNF signaling pathway, MyD88-dependent toll-like receptor signaling pathway, JAK-STAT signaling pathway, regulation of lipopolysaccharide-mediated signaling pathway, and inflammatory bowel disease. Notably, in the comparison between the Rim-treated groups and their vehicle (Veh) counterparts, the Veh-treated group showed greater enrichment in the JAK-STAT signaling pathway, lipopolysaccharide-mediated signaling pathway, and Toll-like receptor signaling pathway.

Subsequently, we performed proteomic analysis using mass spectrometry (MS) to evaluate the impact of Rim on protein expression levels in the hippocampal tissues of 5×FAD mice. With an adjusted *p*-value threshold of < 0.05, approximately 100 differentially expressed proteins (DEPs) were identified in each group when compared to their respective control groups (Figure S3L). Consistent with the transcriptomic findings, the top five KEGG-enriched pathways for DEPs in these groups were predominantly associated with immune-inflammatory processes. These pathways included complement and coagulation cascades as well as staphylococcus aureus infection, identified in the 5×FAD+Veh(F) vs. WT+Veh(F), 5×FAD+Veh(M) vs. WT+Veh(M), and 5×FAD+Rim(F) vs. WT+Veh(F) comparisons. Additionally, the JAK-STAT signaling pathway was enriched in the 5×FAD+Veh(F) vs. WT+Veh(F) group, while Yersinia infection and the chemokine signaling pathway were enriched in the 5×FAD+Veh(M) vs. WT+Veh(M) group (Figure S3M and S3N).

Taken together, these findings suggest that Rim may exert its neuroprotective effects in 7-month-old 5×FAD mice by modulating several immune-inflammatory pathways at both the transcriptional and translational levels.

We next performed weighted gene co-expression network analysis (WGCNA) on all 56,691 genes across 18 samples to further investigate the mechanisms underlying the beneficial effects of Rim in 5×FAD mice. The sample dendrogram and trait heatmap are presented in Figure S4A. Hierarchical clustering identified 27 distinct modules (Figure S4B), with the branches of the dendrogram (meta-modules) grouped based on eigengene correlations (Figure S4C). Among these, three modules— tan, dark grey, and brown—exhibited a robust and statistically significant positive correlation with the 5×FAD group, while being negatively correlated with the WT and Rim-treated groups (Figure S4D and 4A). To refine our analysis, we conducted Mfuzz clustering based on gene expression patterns to identify genes with high expression in Veh-treated 5×FAD mice. This analysis yielded 20 gene expression clusters, of which only Cluster 5 met our criteria: genes highly expressed in the Veh-treated 5×FAD group but attenuated in the Rim-treated 5×FAD group (Figure S4E). To identify hub genes associated with Rim’s effects in AD, we performed an intersection analysis between the three WGCNA modules and Cluster 5, resulting in 683 intersecting genes, which we termed RNA-seq-Rim Pharmaco-genes (RNA-seq-RPG) (Figure 4B). Similarly, we applied the Mfuzz algorithm to soft-cluster proteins and identified two clusters, Cluster 3 and Cluster 20, which we designated as MS-Rim Pharmaco-proteins (MS-RPP) for further investigation (Figure S4F). To narrow down the candidate genes, we intersected the RNA-seq-RPGs with the MS-RPPs, identifying 26 genes that were strongly correlated with Rim’s efficacy at both the transcriptional and translational levels (Figure 4C). A combined heatmap of gene and protein expression showed that all 26 genes followed a consistent trend: they were highly upregulated in Veh-treated 5×FAD mice and attenuated in Rim- treated 5×FAD mice (Figure 4D). Interestingly, nearly half of these genes are known to regulate inflammation (e.g., *Borcs5*, *Hdac11*, *Slc35a2*, *Chtop*, *Hnrnpa2b1*, *Emc3*, *Kcmf1*, *Dgka*, *Dnaja3*, *Smarcb1*, *Wdsub1*), while five others (*Nop56*, *Srsf6*, *Hnmpa1*, *Golm1*, *Srsf2*) have been implicated in AD progression (Figure 4E). Notably, *Hdac11* emerged as a particularly intriguing target. As the sole member of the class IV HDAC subfamily (Yao et al., 2022), HDAC11 has received little attention in the context of AD, despite reports linking other HDAC family members (HDAC3, HDAC5, HDAC6) to the disease (Figure 4E) (Agis-Balboa et al., 2013; Ricciardi et al., 2023; Zhang et al., 2012). Based on these findings, we speculate that HDAC11 may play a critical role in mediating the neuroprotective and anti-inflammatory effects of Rim in 7-month-old 5×FAD mice.

**Figure 4.**
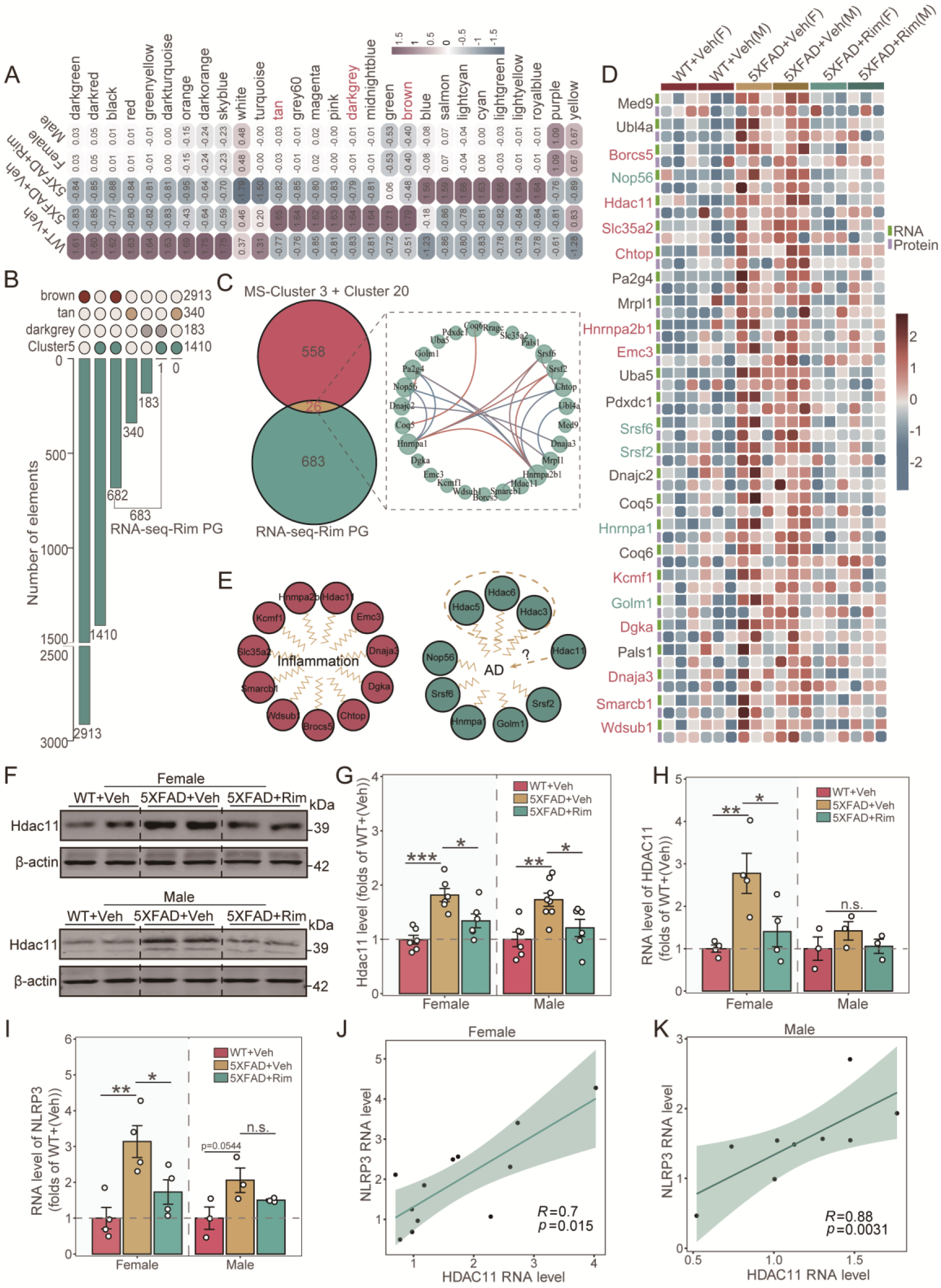
HDAC11 was closely related to the therapeutic efficacy of CGRP antagonist Rim in the hippocampus of 7-month-old 5×FAD mice. (A) WGCNA module trait association correlation graph. (B) WGCNA three AD-related modules and Mfuzz clustering analysis cluster 5 UpSet graph. (C) Venn diagram and protein-protein interaction (PPI) analysis of overlapping sets of RNA-seq- Rim PGs with Cluster 3 and Cluster 20. (D) 26 overlapping genes expression heat map. (E) Classification of inflammation-related genes and AD-related genes. (F and G) Western blot image (F) and densitometry analysis of HDAC11 in the hippocampus (G). (H) Quantitative PCR analysis of *Hdac11* RNA expression level in the hippocampus. (I) Quantitative PCR analysis of *Nlrp3* RNA expression level in the hippocampus. (J and K) Correlation analysis of the RNA levels of *Hdac11* and *Nlrp3* in the hippocampus of female (J) and male mice (K). n=3/genotype. \**p* < 0.05, \*\**p* < 0.01, \*\*\**p* < 0.001, n.s., non-significant. Statistical analyses were analyzed by one-way ANOVA followed by a post hoc Dunnett’s test. See also Figure S4.

We further examined the changes of *Hdac11* expression in the brains of 5×FAD mice and observed a significant upregulation of both transcription and protein levels in the hippocampus of Veh-treated 5×FAD mice compared to WT mice (Figure 4F–4H). Treatment with Rim significantly reversed the protein levels of HDAC11 in the hippocampus of both male and female 5×FAD mice. However, the mRNA expression of *Hdac11* was only significantly reversed in female 5×FAD mice, suggesting a sex-specific effect and highlighting the close association between *Hdac11* and the efficacy of Rim in the hippocampus of 5×FAD mice (Figure 4F–4H). Given that HDAC11 has been reported to promote the NLRP3/caspase-1/GSDMD pathway, leading to pyroptosis via the regulation of ERG acetylation in human umbilical vein endothelial cells (Yao *et al*., 2022), we hypothesize that the neuroprotective effects of Rim in the hippocampus of 5×FAD mice may involve the HDAC11- NLRP3 inflammasome pathway. To validate this hypothesis, we performed qPCR and found that the expression of *Nlrp3* was significantly upregulated in the hippocampus of Veh-treated 5×FAD mice compared to WT mice. Rim treatment significantly downregulated *Nlrp3* mRNA levels in female 5×FAD mice, but not in males (Figure 4I). To further explore the relationship between *Hdac11* and *Nlrp3*, we performed a correlation analysis of their mRNA expression levels in the hippocampus of male and female 5×FAD mice. The results revealed a significant positive correlation between *Hdac11* and *Nlrp3* expression in both sexes (Figure 4J and 4K). Taken together, these findings suggest that HDAC11 may serve as a critical mediator of Rim’s neuroprotective effects in the hippocampus of 5×FAD mice.

### CGRP receptor antagonist Rim reverses alterations in lipid metabolism in 7-month-old 5×FAD mice

To further investigate the impact of Rim on the metabolism of 7-month-old 5×FAD mice, we performed KEGG metabolic pathway enrichment analysis using RNA-seq-RPGs and MS-RPPs. Pathways associated with sugar and lipid metabolism were significantly enriched at both the RNA and protein levels. These included glycerophospholipid metabolism, carbon metabolism, and nucleotide sugar biosynthesis at the transcriptomic level, as well as N-glycan biosynthesis, fatty acid metabolism, and glycerophospholipid metabolism at the proteomic level (Figure 5A and 5B). To distinguish the activity levels of lipid and carbohydrate metabolism in 5×FAD mice, we analyzed the expression patterns of genes involved in these pathways across different experimental groups. The results revealed that lipid metabolism was more active, as genes related to this pathway showed greater dispersion in the ternary plot compared to those involved in carbohydrate metabolism, in both female and male 5×FAD mice (Figure S5A and S5B).

**Figure 5.**
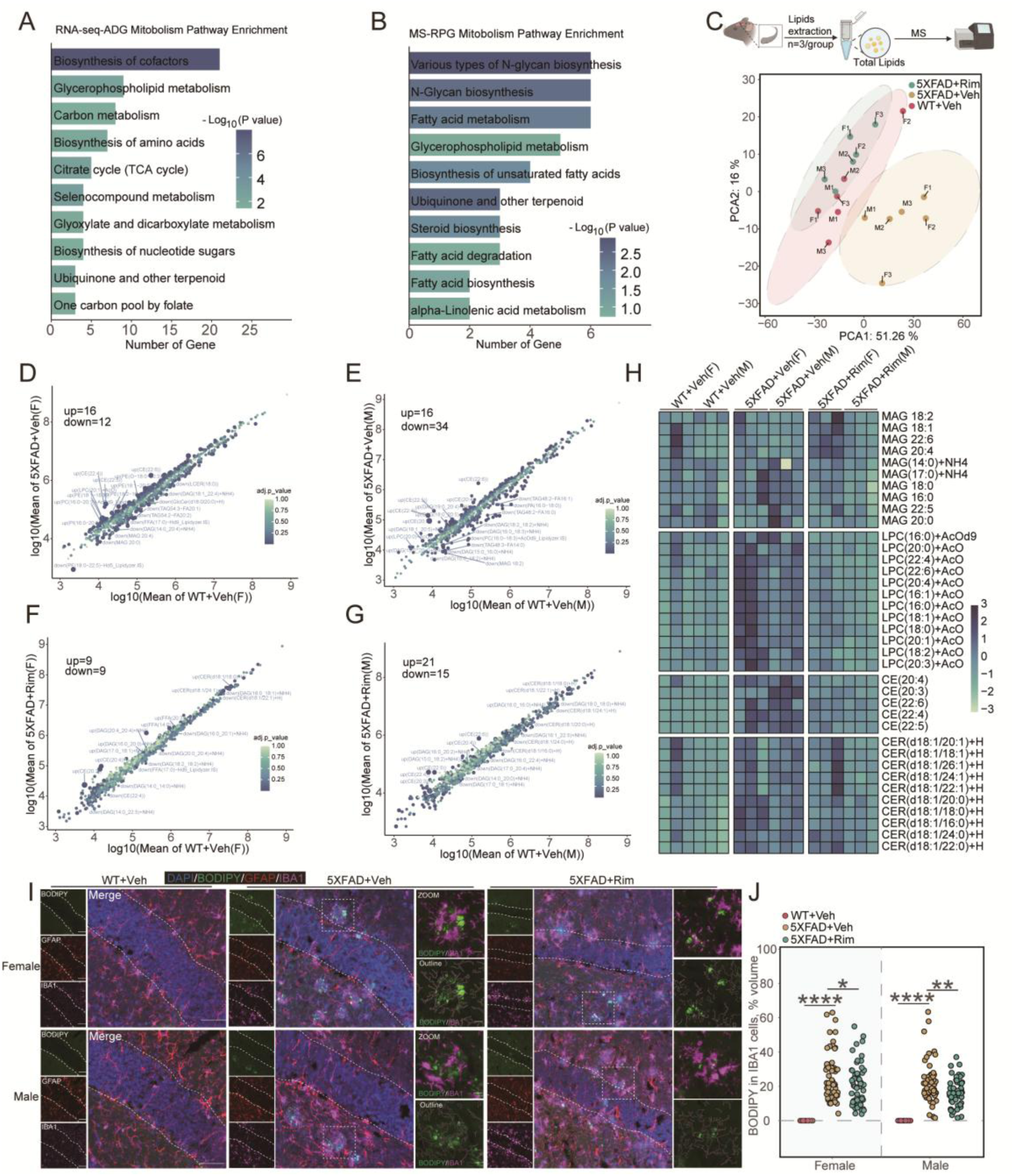
Rim reverses changes in lipid metabolism in 7-month-old 5×FAD mice (A) KEGG metabolism pathways enrichment analysis of RNA-seq-Rim PGs. (B) KEGG metabolism pathways enrichment analysis of Cluster 3 and Cluster 20. (C) Top, Schematic representation of the experimental design; bottom, PCA based on results of lipidomic profiles obtained from MS. (D) Volcano plot of differentially expressed lipids (DELs) in 5×FAD+Veh(F) vs. WT+Veh(F) mice. (E) Volcano plot of DELs in 5×FAD+Veh(M) vs. WT+Veh(M) mice. (F) Volcano plot of DELs in 5×FAD+Rim(F) vs. WT+Veh(F) mice. (G) Volcano plot of DELs in 5×FAD+Rim(M) vs. WT+Veh(M) mice. (H) Monoacylglycerol (MAG), lysophosphatidylcholine (LPC), cholesteryl ester (CE) and ceramide (CER) expression heat map. (I and J) Representative BODIPY (green), GFAP (red) and IBA-1 (pink) co-staining(I) and quantification (J) in the hippocampus of 7-month-old mice. n=60 IBA-1^+^ cell, 6/group. Scale bars: 50 μm and 10 μm (ZOOM) in (I). \**p* < 0.05, \*\**p* < 0.01, \*\*\*\**p* < 0.0001. Statistical analyses were analyzed by one-way ANOVA followed by a post hoc Dunnett’s test. See also Figure S5.

To identify specific lipid species dysregulated in 5×FAD mice, we conducted lipidomic profiling of hippocampal tissues. PCA analysis revealed that the principal components of 5×FAD-Veh mice were significantly separated from the Veh-WT group, whereas 5×FAD-Rim mice were not significantly separated from Veh-WT mice (Figure 5C). Among the 700+ lipid species measured in the hippocampus, we identified 28 differentially expressed lipids (DELs) in 5×FAD+Veh(F) vs. WT+Veh(F) (Figure 5D), 50 DELs in 5×FAD+Veh(M) vs. WT+Veh(M) (Figure 5E), 18 DELs in 5×FAD+Rim(F) vs. WT+Veh(F) (Figure 5F), and 36 DELs in 5×FAD+Rim(M) vs. WT+Veh(M) (Figure 5G). Notably, among the decreased lipids were triacylglycerol (TAG) and diacylglycerol (DAG) species, while cholesteryl ester (CE) and lysophosphatidylcholine (LPC) species were significantly increased. These findings highlight specific disruptions in cholesterol and fatty acid metabolism in 7-month-old 5×FAD mice (Figure 5D–5G). Emerging research in AD has increasingly focused on identifying lipid signatures associated with neuroinflammation and pathologies, including alterations in monoacylglycerol (MAG), LPC, CE, and ceramide (CER) species (Ferré-González et al., 2024; Litvinchuk et al., 2024). When comparing the cumulative effects on hippocampal lipidomes between WT, Veh-5×FAD, and Rim-5×FAD mice, we observed significant elevations in LPC, CE, and CER levels in Veh-5×FAD mice (Figure 5H). Remarkably, treatment with Rim substantially mitigated these alterations (Figure 5H). To further investigate the effect of Rim on lipids accumulation, we performed BODIPY staining in the hippocampus and observed a significant reduction in BODIPY signals especially in microglia of Rim-treated 5×FAD mice compared to 5×FAD-Veh mice (Figure 5I and 5J). These results suggest that Rim may synergistically rescue impaired cholesterol metabolism, improve endolysosomal lipid clearance, and reduce neuroinflammation in 7-month-old 5×FAD mice.

### HDAC11 inhibition mitigates neuronal cell death induced by sAβ1-42 *in vitro* and alleviates paralysis in *Caenorhabditis elegans* CL2006

To investigate the cell-specific expression of Hdac11 in AD, we retrieved snRNA-Seq data from the SEA-AD database (Gabitto *et al*., 2024). As shown in Figure S6A–S6D, Hdac11 is widely expressed across various brain cell types, including neurons, microglia, and astrocytes. We then evaluated the effects of CGRP receptor antagonist Rim on HDAC11 level in primary neurons and mixglia treated with 1 μM Rim with or without 10 mM sAβ1-42 for 24 h (Figure S6E). Our analysis revealed a significant increase in HDAC11 expression in both primary neurons and mixed glial cells following treatment with sAβ1-42 (Figure S6F- S6H). Although 24 h of Rim treatment did not markedly decrease HDAC11 expression in mixglia, we observed a significant reduction of HDAC11 levels in primary neurons treated with Rim (Figure S6F- S6H). These findings suggest that Rim exerts its neuroprotective effects predominantly by directly modulating HDAC11 activity in neurons rather than glial cells. To further explore the role of HDAC11 in AD pathology, we examined the effects of the HDAC11 inhibitor Elevenostate (Ele) on neuronal responses to sAβ1-42-induced injury. Primary cortical neurons from WT mice were treated with 10 mM sAβ1-42 in the presence or absence of Ele for 24 h (Figure S6I). The results of CCK8 assay and Hoechst 33342/PI staining indicated that Ele (10 μM) significantly improved cell viability compared to the sAβ1-42-treated group (Figure S6J- S6L). Additionally, Ele at concentrations of 4 μM and 20 μM was found to alleviate paralysis in the Aβ1-42 transgenic *Caenorhabditis elegans* (CL2006) model of AD (Figure S6N).

### Short-term inhibition of hippocampal HDAC11 function reduced astroglial activation in female 5×FAD mice

To determine whether HDAC11 inhibition contributes to the protective effects of Rim on AD pathology and neuroinflammation *in vivo*, we performed bilateral stereotaxic injections of Ele into the CA1 region of the hippocampus in 7-month-old 5×FAD mice (Figure S7A). One day after Ele treatment, memory and anxiety-like behaviors were assessed using the Y-maze and EPM tests. As shown in Figure S7B- S7D, this short-term Ele treatment did not induce significant changes in anxiety-like behavior or memory deficits in 5×FAD mice. Thioflavin-S staining revealed no significant differences in Aβ plaque burden in the hippocampus of Ele-treated 5×FAD mice compared to the control group (Figure S7E- S7G). Among other AD pathological markers, Ele treatment significantly reduced both the number of astrocytes and the expression level of GFAP proteins in the hippocampus of female 5×FAD mice (Figure S7H- S7I). However, Ele did not have a significant impact on tau pathology, the number of microglia, or the expression levels of HDAC11, APP, and IBA-1 (Figure S7K- S7M). When assessing apoptotic protein levels in the hippocampus, we observed that Ele treatment significantly upregulated the expression level of Bcl-2 and decreased the Bax/Bcl-2 ratio only in the hippocampus of female 5×FAD mice compared to the control group (Figure S7K–S7L). These findings suggest that Ele may confer a protective effect on the hippocampus of 7-month-old female 5×FAD mice, potentially through mechanisms involving astrocytic modulation rather than direct effects on Aβ pathology, tau pathology, or microglial activity.

### *Hdac11* knockdown in the hippocampus of 7-month-old 5×FAD mice results in enhanced memory

The Hdac11 inhibitor Ele used in this study was neither a long-term therapeutic intervention nor specifically targeted to mouse brain neurons, which may have contributed to its limited effects on the AD pathology described above. To address these limitations, we employed recombinant adeno-associated viral vectors (rAAV) to achieve RNAi-mediated *Hdac11* knockdown specifically in the hippocampus in 5×FAD mice. rAAV vectors containing either hSYN-EGFP-shRNA-Hdac11 (shRNA-Hdac11) or hSYN-EGFP (shRNA-Control) were bilaterally injected into the hippocampal CA1 regions of 7-month-old 5×FAD mice (Figure 6A). One-month after viral injection, EGFP expression was observed broadly in the hippocampal pyramidal fields (including CA1, CA2, CA3, DG) of both groups (Figure 6B). Immunofluorescence analysis revealed that most EGFP-positive cells were co-labeled with NeuN (a neuronal marker) but not with IBA-1 (a microglial marker) or GFAP (an astrocyte marker), confirming that the virus predominantly infected neurons (Figure 6C and 6D). Furthermore, protein analysis validated that Hdac11 expression was significantly reduced in the hippocampus of rAAV shRNA-Hdac11-injected mice compared to the control mice (Figure 6E and 6F). Anxiety-related behavioral assessments were conducted using the EPM and FST paradigms. We observed that *Hdac11* knockdown significantly reduced the anxiety index in female 5×FAD mice, as assessed by the EPM, but no significant changes were noted in the FST (Figure 6G-6I). Memory related behavioral assessments were performed to evaluate the effect of *Hdac11* knockdown on the hippocampus-dependent spatial learning and memory. 5×FAD mice injected with rAAV shRNA-Hdac11 exhibited significant enhancements in the NLR test (Figure 6J), Y-maze test (Figure 6L and 6M), and MWM test (Figure 6N–6R). Specifically, rAAV shRNA-Hdac11-injected female mice recovered their ability in the NLR test (Figure 6J) and reversed alternation deficits in the Y-maze test. Male AD mice also showed improved alternation performance in the Y-maze test (Figure 6L and 6M). In the MWM test, strikingly, female mice infected with rAAV shRNA-Hdac11 exhibited significantly shorter latencies to locate the platform on day 6 of training (Figure 6N–6R). During the probe trial, 5×FAD mice injected with rAAV shRNA-Hdac11 demonstrated superior performance. Specifically, mice injected with rAAV shRNA-Hdac11 spent more time in the platform quadrant, and female mice also exhibited more platform crossings compared to their control counterparts (Figure 6N–6R). Collectively, these findings demonstrate that *Hdac11* knockdown in the hippocampus of 5×FAD mice was sufficient to enhance hippocampus-dependent learning and memory.

**Figure 6.**
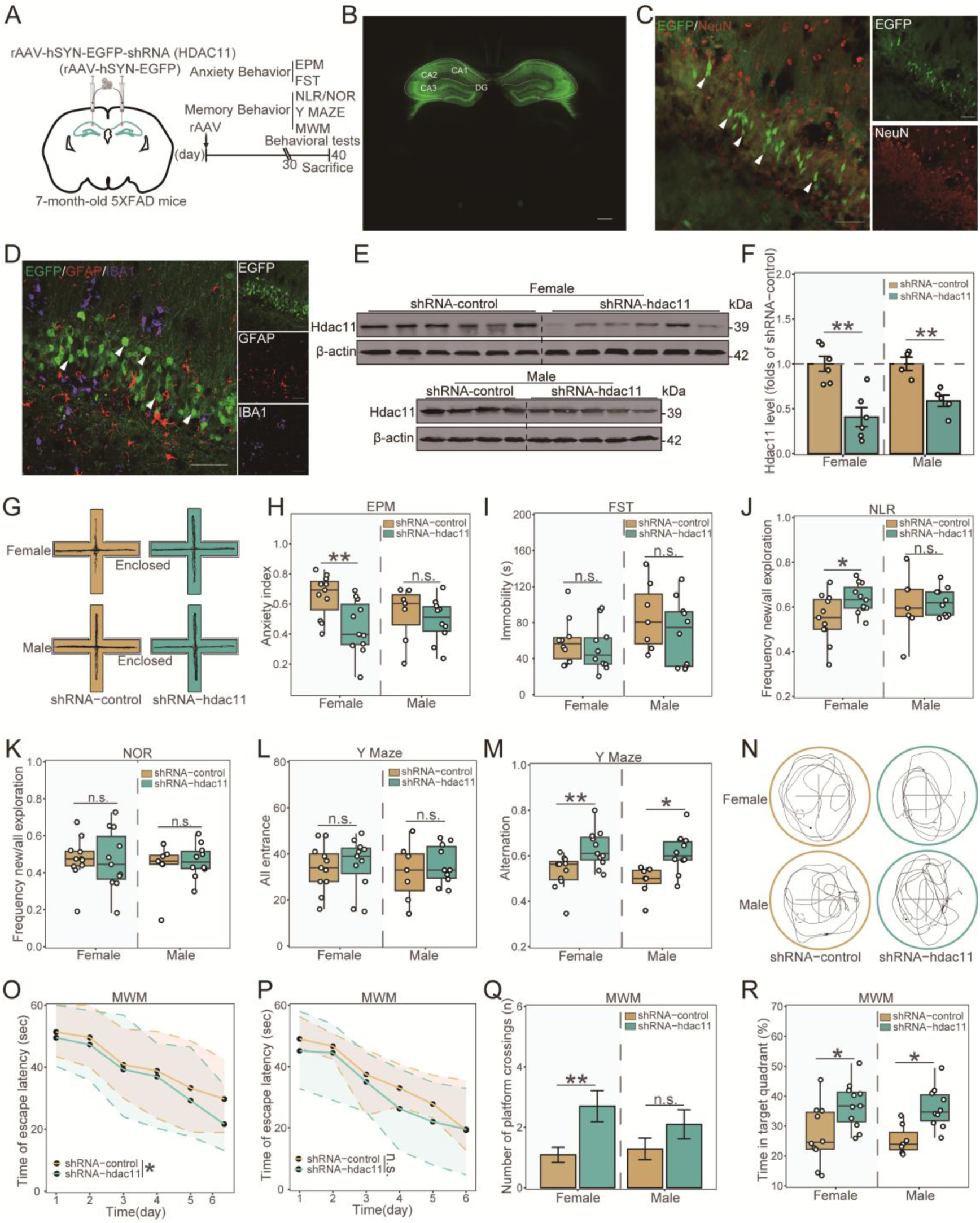
*Hdac11* knockdown in the hippocampus of 7-month-old 5×FAD mice results in enhanced memory (A) Schematic representation of the experimental design. (B) Brain slices stained with EGFP (viral infection). (C) Brain slices were stained with EGFP (viral infection) and NeuN (neurons) antibodies. (D) Brain slices were stained with EGFP (viral infection), GFAP (astrocyte) and IBA-1 (microglia) antibodies. (E and F) Western blot image (E) and densitometry analysis of Hdac11 in the hippocampus (F). (G) Representative movement traces of the mice in the Elevated plus maze (EPM). (H) The anxiety index calculated by the EPM. (I) Immobility time in the Forced swim test (FST). (J) The ratio of the number of explorations of the new location object to the total number of explorations in the Novel location recognition (NLR) task. (K) The ratio of the number of explorations of the new object to the total number of explorations in the Novel object recognition (NOR) task. (L and M) The number of total arm entries (L) and spontaneous alternations (M) in the Y-maze test. (N) Combined traces of the mice from each group during the probe test in the Morris water maze (MWM). (O-R) The results of Morris water maze (MWM). (O, P) Latency to the platform in female (O) and male (P); (Q)Entries into the platform zone; (R) The percentage of time spent in target quadrant. in the MWM. n=7-11/genotype. Scale bars: 1mm in (B); 50 μm in (C and D). \**p* < 0.05, \*\**p* < 0.01, n.s., non-significant. Statistical analyses were analyzed by unpaired Students’ t-test.

### AD-related pathology in the hippocampus of 7-month-old 5×FAD mice would be alleviated by knockdown of *Hdac11*

We further investigated AD-associated pathological changes in *Hdac11* knockdown 5×FAD mice. Thioflavin-S staining revealed a significant reduction in both the number and size of Aβ plaques in the hippocampus of 5×FAD mice injected with rAAV shRNA-Hdac11 compared to mice in the control group (Figure 7A–7C). Additionally, rAAV shRNA-Hdac11-infected mice exhibited a dramatic decrease in the astrocyte number and GFAP protein levels in the hippocampus (Figure 7D and 7E). Interestingly, while a reduction in microglial cell number was observed only in rAAV shRNA-Hdac11-treated female 5×FAD mice, both male and female AD mice with *Hdac11* knockdown showed a significantly reduced IBA-1 protein expression (Figure 7D-7I). Moreover, 5×FAD mice infected with rAAV shRNA-Hdac11 demonstrated a greater than 50% reduction in APP levels and a significant decrease in phosphorylated tau proteins in the hippocampus (Figure 7G-7I). To further explore the effects of HDAC11 inhibition on cell survival, we examined the expression of Bcl-2 and Bax in the hippocampus of 5×FAD mice. Compared to the Control group, rAAV shRNA-Hdac11-injected mice displayed a significantly lower Bax expression and a markedly reduced Bax/Bcl-2 ratio, particularly in female mice (Figure 7G-7J). Finally, we performed BODIPY staining to assess lipid accumulation in the hippocampus of rAAV shRNA-Hdac11- injected mice and observed a significant reduction in BODIPY signal intensity in microglia compared to Control mice (Figure 7K and 7L). In summary, these results demonstrate that *Hdac11* knockdown in hippocampal neurons is sufficient to alleviate the core pathological hallmarks associated with AD, including Aβ accumulation, neuronal death, neuroinflammation, tau hyperphosphorylation, and excessive lipid accumulation.

**Figure 7.**
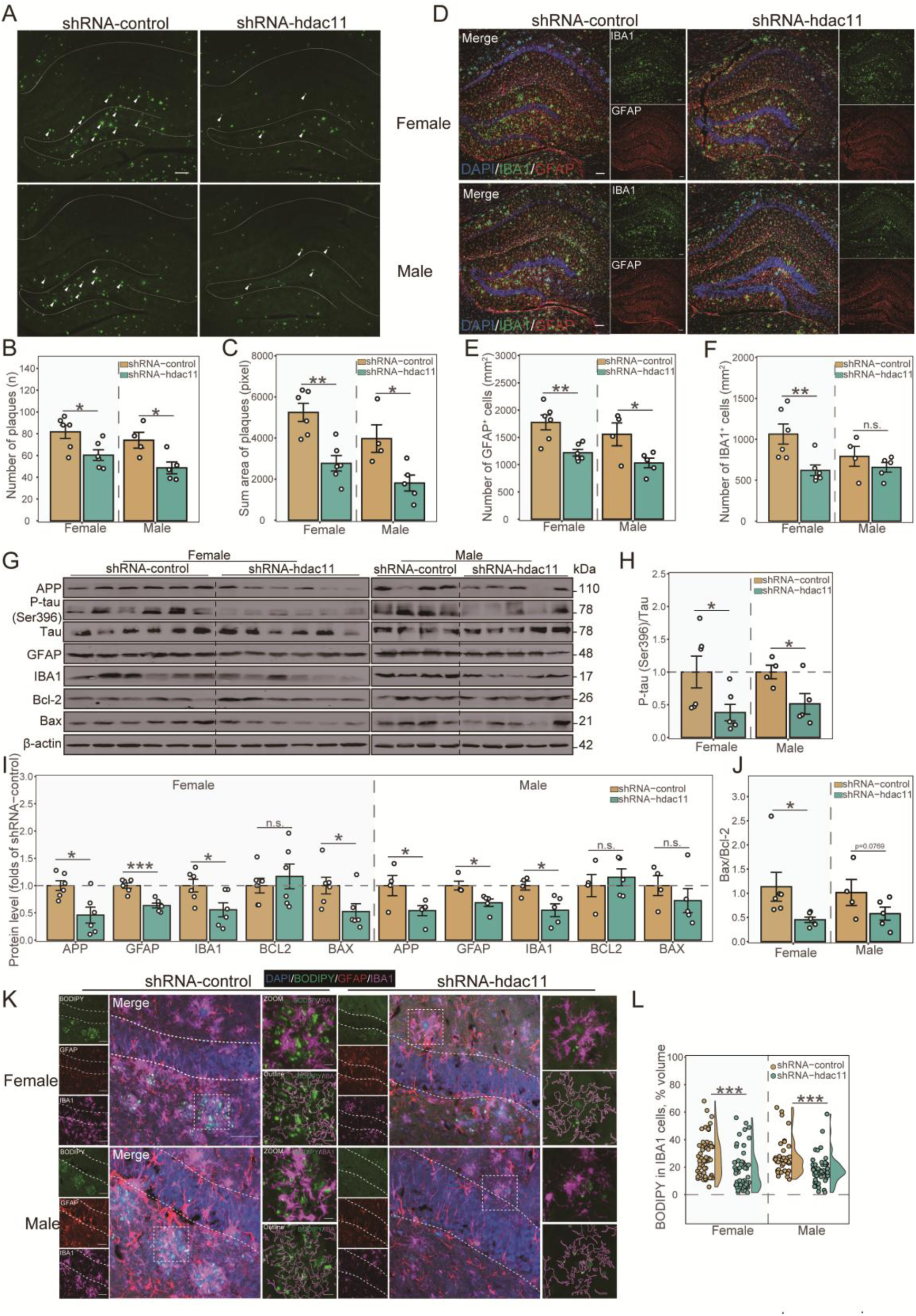
AD-related pathology in the hippocampus of 7-month-old 5×FAD mice would be alleviated knockdown of *Hdac11*. (A) Representative Thioflavin-S staining hippocampal images of 5×FAD mice injected with rAAV expressing control shRNA or HDAC11 shRNA. (B and C) Quantitative measurements of Aβ plaque numbers (B) and area (C) in the hippocampus. (D) Immunofluorescence staining of GFAP (red) and IBA-1 (green)in the hippocampus. (E and F) Quantifications of GFAP^+^ astrocytes (E) and IBA-1^+^ microglia (F) in the hippocampus. (G-J) Western blot image (G) and densitometry analysis of APP, phosphorylated tau, total tau, GFAP, IBA-1, Bcl-2 and Bax in the hippocampus (H-J). (K and L) Representative BODIPY (green), GFAP (red) and IBA-1 (pink) co-staining (K) and quantification (L) in the hippocampus of 7-month-old mice. n=4-6/genotype (B-J); n=30 IBA-1^+^ cell, 4-6/group (L). Scale bars: 100 μm in (A and B); 50 μm and 10 μm (ZOOM) in (K). \**p* < 0.05, \*\**p* < 0.01, \*\*\**p* < 0.001, n.s., non-significant. Statistical analyses were analyzed by unpaired Students’ t-test.

### Knockdown of *Hdac11* exerts a protective role in AD by targeting LXRβ/ABCA1 axis

To investigate the potential regulatory mechanisms of HDAC11, we isolated hippocampal lysis from WT mice and performed immunoprecipitation with HDAC11 antibody or IgG followed by mass spectrometry (IP-MS) analysis (Figure 8A). Correlation analysis of enriched proteins revealed a higher degree of similarity between samples within the same antibody group compared to samples from different antibody group (Figure 8B). After excluding proteins that interacted with IgG, which were defined as non-specific bindings, we identified a total of 154 highly specific HDAC11- interacting proteins (Figure 8C). Subsequent KEGG pathway annotation classified these 154 HDAC11-interacting proteins into pathways associated with neurodegenerative diseases, inflammation, and lipid metabolism (Figure 8D). In parallel, hippocampal lysates from WT and *Calca* knockout (KO) mice were immunoprecipitated using acetylated lysine antibody to identify highly acetylated proteins that may serve as potential HDAC11 targets. Correlation analysis again showed a higher degree of similarity between samples within the same genotype group compared to samples from different genotype group (Figure 8E). Compared to the WT group, we identified 74 highly acetylated proteins in the *Calca* KO group (Figure 8F). To further refine the list of potential functional Hdac11 target proteins, we intersected the 74 highly acetylated proteins from the *Calca* KO group with the 154 HDAC11-interacting proteins identified via IP-MS (Figure 8C). This analysis yielded 27 candidate proteins for further investigation (Figure 8G and 8H). Among these candidates, the protein of primary interest is NR1H2 (also known as LXRβ), due to its critical role in promoting the expression of ABCA1. Enhanced ABCA1 expression has been shown to improve cognitive function in multiple AD mouse models by regulating lipid metabolism and exerting potent anti-inflammatory effects through the suppression of pro-inflammatory gene expression (Kang and Rivest, 2012; Lewandowski et al., 2022). Notably, an evolutionary conservation analysis of NR1H2/LXRβ across four species revealed that three lysine residues near the N-terminal and C-terminal regions are highly conserved, suggesting these lysine residues may perform an evolutionarily conserved function (Figure 8I). Therefore, NR1H2/LXRβ may represent a key target through which HDAC11 inhibition exerts its neuroprotective and anti-inflammatory effects in 5×FAD mice.

**Figure 8.**
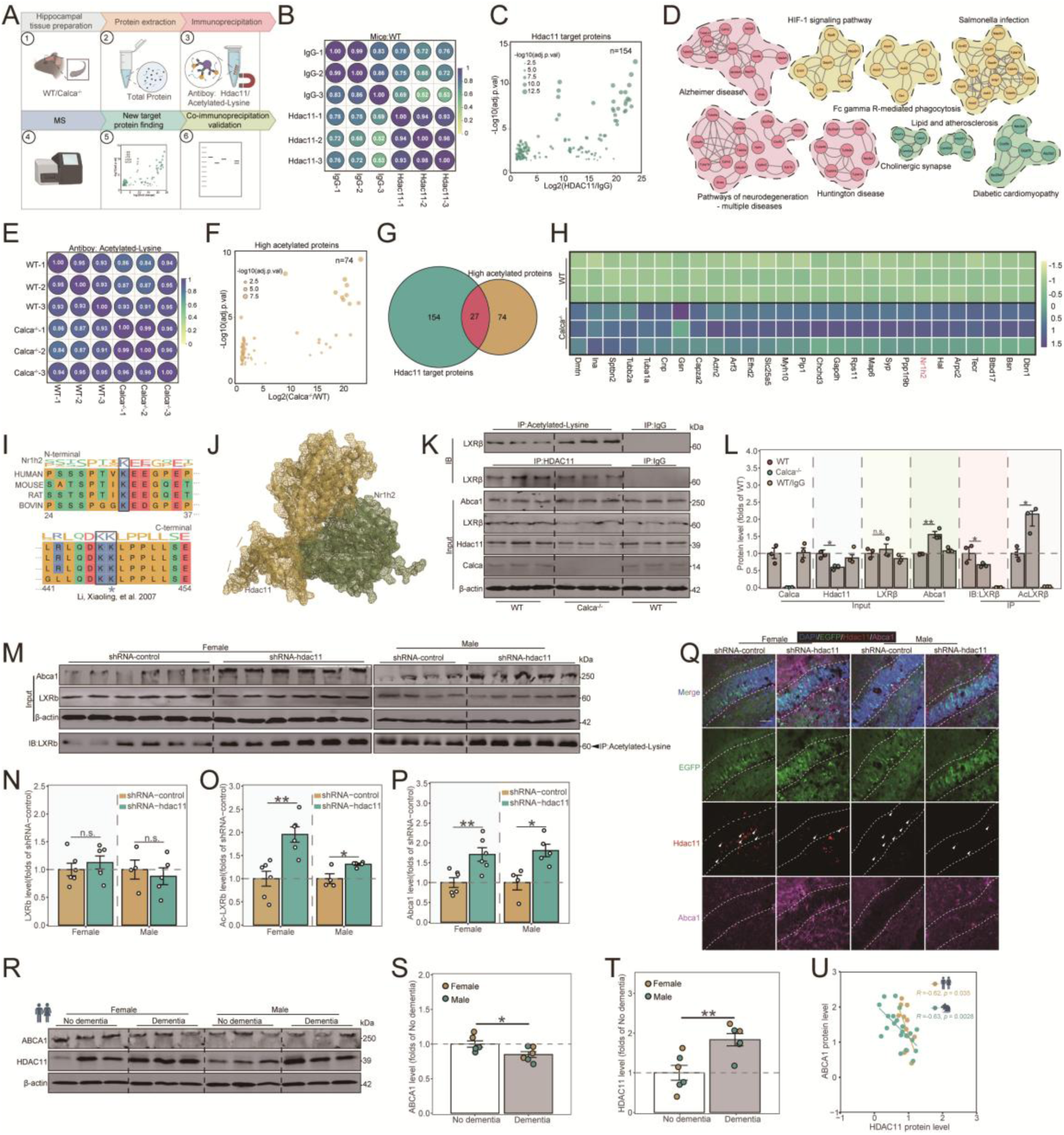
Knockdown of *Hdac11* exerts a protective role *in vivo* by targeting LXRβ/ABCA1 axis. (A) Schematic representation of the experimental design. (B) Correlation analysis of IgG and Hdac11 antibody groups. (C) Scatter plot depicting the 154 HDAC11-interacting candidates from the immunoprecipitation/mass spectrometry (IP-MS). (D) Protein-protein interaction (PPI) network analysis of the HDAC11 target proteins that belong to the enriched KEGG pathways. (E) Correlation analysis of WT and *Calca^-/-^* groups. (F) Scatter plot depicting the 74 highly acetylated proteins from the IP-MS. (G) Venn diagram of overlapping sets of Hdac11 target proteins with highly acetylate proteins. (H) 27 overlapping proteins expression heat map. (I) Protein sequence alignment of NR1H2/LXRβ across species. (J) Protein–protein docking and molecular dynamics of HDAC11 (yellow) bound to NR1H2 (green). (K and L) Hippocampal protein lysates from WT or *Calca^-/-^*mice were precipitated with HDAC11, acetylated-lysine, or IgG antibodies and then analyzed by Western blotting (K), followed by statistical analysis (L). (M-P) Hippocampal protein lysates from shRNA-control or shRNA-Hdac11 administrated 5×FAD mice were precipitated with acetylated-lysine antibodies and then analyzed by Western blotting (M) followed by statistical analysis (N-P). (Q) Brain slices were stained with EGFP (green), HDAC11 (red), and ABCA1 (pink) antibodies. Scale bar= 50 μm. (R-T) Western blot image (R) and densitometry analysis of HDAC11 (S) and ABCA1 (T) in human hippocampus. (U) Correlation analysis of the protein levels of HDAC11 and ABCA1 in human and mice hippocampus. n (WT)=3; n (*Calca^-/-^*)=3; n (shRNA-control)=4-6; n (shRNA-Hdac11)=4-6. n (No dementia)=6; n (Dementia)=6. Scale bars: 50 μm in (Q). \**p* < 0.05, \*\**p* < 0.01, n.s., non-significant. Statistical analyses were analyzed by unpaired Students’ t-test. The Spearman method was used for the correlation analysis (U). See also Figure S8.

To further explore this interaction, we performed molecular docking of HDAC11 with NR1H2/LXRβ. This analysis identified 10 binding modes between HDAC11 and LXRβ, with confidence scores ranging from 0.8634 to 0.9569 and docking scores spanning from -242.19 to 304.95 (Figure S8A). The best docking results obtained from the HDOCK server are shown in Figure 8J. These results confirmed that HDAC11 has a high affinity with the target protein LXRβ. To determine whether the CALCRL/HDAC11/LXRβ/ABCA1 pathway functions *in vivo*, protein levels of this axis were initially assessed in the hippocampus of mice. As shown in Figure 8K and 8L, HDAC11 protein expression was significantly reduced in *Calca^−/−^* group compared to WT mice. Conversely, *Calca* knockout significantly increased ABCA1 protein expression, while having no significant effect on LXRβ protein levels (Figure 8K and 8L). To further examine whether ABCA1 is regulated by HDAC11, immunoprecipitation was performed using HDAC11 antibodies. The results demonstrated that *Calca* knockout disrupted the formation of HDAC11-LXRβ complexes. Additionally, to evaluate whether HDAC11 modulates the acetylation status of LXRβ in mouse brain, protein lysates from the hippocampus were immunoprecipitated with acetylated-lysine antibodies and subsequently analyzed by Western blotting. The findings revealed that *Calca* knockout markedly increased the acetylation of LXRβ compared to the WT group. Based on these results, we infer that HDAC11 mediates its effects on AD pathology through the LXRβ/ABCA1 axis.

To verify that HDAC11 is a regulating molecule of the LXRβ/ABCA1 axis in AD pathology, we cultured hippocampal neurons from WT and *Calca* knockout mice, and treated them with 10 μM sAβ1-42 for 24 h. As shown in Figure S8B and S8C, HDAC11 expression was reduced by approximately 50% in the *Calca* knockout group compared to the WT group, regardless of sAβ1- 42 treatment. Meanwhile, the results of Western blotting and immunofluorescence confirmed an increase in ABCA1 protein expression and a decrease in the BODIPY^+^ dot area in the *Calca* knockout group compared to the WT group, irrespective of sAβ1-42 treatment (Figure S8B-S8I). To further validate these findings *in vivo*, we performed immunoprecipitation on hippocampal protein lysates from 5×FAD mice infected with rAAV shRNA-Hdac11 or rAAV-GFP, using acetylated-lysine antibodies. The results showed that *Hdac11* knockdown did not alter the expression level of LXRβ but significantly increased its acetylation, which in turn elevated ABCA1 protein levels compared to the control group (Figure 8M-8P). These findings were further corroborated by immunofluorescence staining of HDAC11 and ABCA1 in mouse hippocampus (Figure 8Q). Interestingly, following sAβ1-42 treatment, Western blot and ELISA analyses revealed no significant increase in Calca levels in WT neuronal lysates (Figure S8B-S8E). In contrast, a marked elevation of CALCA concentration in the supernatant of neuronal cultures was detected by ELISA (Figure S8B-S8D). These findings suggest that during the pathological progression of AD, the accumulation of Aβ in neurons may accelerate the release of CALCA, which then inhibits LXRβ acetylation via activation of the CALCRL/HDAC11 pathway. Such inhibition of LXRβ acetylation ultimately downregulates ABCA1 expression, contributing to lipid metabolism dysregulation. To evaluate whether the HDAC11/LXRβ/ABCA1 axis is relevant to humans, we performed Western blot analysis on post-mortem brain samples from individuals with or without dementia. The results demonstrated a significant increase in HDAC11 expression in dementia patients compared to non-dementia controls, accompanied by a marked decrease in ABCA1 expression (Figure 8R-8T). Correlation analysis of hippocampal HDAC11 and ABCA1 protein expression levels in both 5×FAD mice and humans revealed a significant negative correlation between these two proteins in both species (Figure 8U). These findings suggest that HDAC11 may modulate AD pathology through the LXRβ/ABCA1 axis in mice and humans.

## DISCUSSION

In the present study, we observed that the CGRP receptor component CALCRL was highly upregulated in the hippocampal tissues of both AD dementia patients and the 5×FAD mice. Furthermore, inhibition of CGRP receptor function, achieved through ligand knockout (*Calca^-/-^*) or using the receptor inhibitor Rimegepant (Rim), attenuated sAβ1-42-induced neuronal death in primary cultures and reduced inflammatory responses in primary glial cells *in vitro*. We further investigated the potential therapeutic effects of CGRP receptor inhibition by utilizing Rim, a small molecule with good blood-brain barrier permeability *in vivo.* Sex is known to influence the onset and progression of Alzheimer’s disease (Pike, 2017), and prior studies have demonstrated that female 5×FAD mice display an earlier onset and more severe AD pathology compared to their male counterparts (Bouter et al., 2021). To avoid confounding effects of sex-specific differences, we examined the effects of Rim in both male and female 5×FAD mice at 7 months of age. Consistent with previous reports, we found that female 5×FAD mice showed more severe AD pathology. Notably, treatment with Rim improved anxiety-like behaviors and spatial memory deficits in both male and female 5×FAD mice in multiple behavioral paradigms.

Memory deficits in AD patients are predominantly attributed to aggressive amyloid plaque deposition, neurofibrillary tangles caused by tau hyperphosphorylation, and reduced synaptic plasticity (Xu *et al*., 2021). In our study, Thioflavin-S staining and Western blot analyses revealed that Rim treatment alleviated amyloid plaque pathology in the hippocampus of 5×FAD mice compared to the Veh-treated group, although no significant differences were observed in the cortex. Similarly, Rim significantly reduced tau phosphorylation levels in the hippocampus but not in the cortex of 5×FAD mice. An increase in synaptic protein levels (SYN) was observed exclusively in the hippocampus of Rim-treated female 5×FAD mice. Excessive neuronal death is another critical contributor to memory deficits in AD patients (Lee et al., 2023). In our study, Rim treatment significantly reduced the number of FJB-positive cells in female 5×FAD mice, as well as the Bax/Bcl-2 ratio in the hippocampus of 5×FAD mice. Neuroinflammation, a key driver of AD pathogenesis, is primarily mediated by activated microglia and astrocytes (Leng and Edison, 2021). In 5×FAD mice, the expression levels of glial marker proteins GFAP and IBA-1 were markedly elevated in both the hippocampus and cortex. However, Rim treatment significantly reduced astrocytic and microglial activation in these regions. Additionally, the pro-inflammatory cytokine IL-1β, which was highly expressed in Veh-treated 5×FAD mice, was significantly decreased following Rim treatment. Taken together, our findings reveal regional heterogeneity in AD pathology in 5×FAD mice and the therapeutic effects of Rim. Specifically, the impact of Rim on amyloid plaque burden, tau phosphorylation, neuronal death, and neuroinflammation was predominantly observed in the hippocampus of 5×FAD mice.

To investigate the potential mechanisms by which Rim alleviates AD pathology, we performed bulk RNA-seq and protein MS analyses on the hippocampus of 5×FAD mice. Using DESeq2, we conducted differential expression analysis between AD and WT groups to derive an AD- and Rim- related signatures. Those signatures included DEGs and DEPs associated with pathways involved in inflammation and immune activation. To further refine our analysis, we applied WGCNA and Mfuzz clustering to the RNA-seq data, identifying 683 Rim-regulated potential genes (RNA-seq- Rim PGs). By intersecting these with 588 Rim-regulated potential proteins (MS-RPPs) identified through Mfuzz clustering of the proteomic data, we pinpointed 26 genes that were highly correlated with the efficacy of Rim at both the transcriptomic and proteomic levels in 5×FAD mice. Notably, over half of these 26 genes are known to regulate inflammation and AD progression, and the one we are most concerned about is Hdac11, the sole member of the class IV HDAC subfamily. HDAC11 has been implicated in promoting the NLRP3/Caspase-1/GSDMD pathway, which leads to pyroptosis, as well as the NLRP3/Caspase-1/IL-1β pathway, which drives inflammation (Yao *et al*., 2022). While HDAC3, HDAC5, and HDAC6 have been reported to play significant roles in AD pathogenesis (Agis-Balboa *et al*., 2013; Ricciardi *et al*., 2023; Zhang *et al*., 2012), there is currently no evidence linking HDAC11 to AD. In our study, we found that the expression of HDAC11 was elevated in the hippocampus of 5×FAD mice, but such increase was reversed by Rim treatment. Moreover, HDAC11 expression was positively correlated with NLRP3 at the transcriptional level, suggesting that the therapeutic efficacy of Rim in AD may, at least in part, be mediated by the inhibition of HDAC11. In addition, KEGG enrichment analysis revealed that lipid metabolism pathways were highly enriched in both RNA-seq-Rim PGs and MS-RPPs datasets. Interactions between lipid metabolism and AD pathogenesis have been extensively studied (Chen et al., 2024; Ferré-González *et al*., 2024; Litvinchuk *et al*., 2024). Using untargeted lipid metabolomics, we observed significant elevations in the levels of LPC, CE, and CER in the hippocampus of 5×FAD- Veh mice. However, these alterations were mitigated by Rim treatment, suggesting that Rim may exert its therapeutic effects by synergistically improving impaired cholesterol metabolism, enhancing endo-lysosomal lipid clearance, and reducing subsequent neuroinflammation.

To investigate whether Rim-mediated Hdac11 inhibition is cell-type specific in the context of AD, we first validated that Rim suppresses the expression of *Hdac11* induced by sAβ1-42 in primary neurons. Furthermore, the selective HDAC11 inhibitor Ele was shown to reduce Aβ-induced neuronal death and paralysis in CL2006 nematodes. Inhibition of HDAC11 for one month in hippocampal neurons of 5×FAD mice significantly alleviated cognitive deficits and AD-related pathologies, including a reduction in Aβ plaque area and number, decreased activation of astrocyte and microglia, and lower levels of APP and phosphorylated tau proteins. Additionally, HDAC11 inhibition decreased Bax expression and the Bax/Bcl-2 ratio in the female hippocampus, indicating an attenuation of pro-apoptotic signaling. HDAC11 inhibition also improved lipid metabolism by reducing lipid accumulation in microglia, further highlighting its potential in mitigating AD-related neuroinflammation and metabolic dysregulation or dysfunction.

To investigate the regulatory mechanism of HDAC11 in AD, we identified 154 HDAC11-interacting proteins using immunoprecipitation-mass spectrometry (IP-MS). Functional enrichment analysis of these interacting proteins suggested that HDAC11 may play a role in the regulation of neurodegenerative diseases, inflammation, and lipid metabolism. Further comparison with highly acetylated proteins in *Calca* KO mice identified 27 potential HDAC11 targets, including NR1H2 (also known as LXRβ), a protein reported to regulate lipid metabolism and inflammation via Abca1, thereby improving cognitive function in AD models (Kang and Rivest, 2012; Lewandowski et al., 2022).

In *Calca* KO mice, HDAC11 expression was reduced while ABCA1 expression was elevated. These mice also exhibited decreased formation of HDAC11-LXRβ complexes and enhanced acetylation of LXRβ. Consistent with these findings, HDAC11 inhibition in 5×FAD mice led to increased ABCA1 levels and LXRβ acetylation, further supporting the LXRβ/ABCA1 axis in mitigating AD pathology. Analysis of human post-mortem brain tissue revealed a negative correlation between HDAC11 and ABCA1 expression, suggesting that HDAC11 regulates AD pathology through the LXRβ/ABCA1 axis in both mouse models and humans.

It is well known that CGRP and its receptor are endogenous mediators of migraines, playing a critical role in migraine pathophysiology (Ho *et al*., 2010). The release of CGRP facilitates vasodilation and neurogenic inflammation, key processes in the development of migraines (Malhotra, 2016). Inhibition of the CGRP receptor using monoclonal antibodies and antagonists has demonstrated efficacy in reducing both the frequency and severity of migraines, with clinical trials supporting its use as an effective acute treatment (Dodick, 2019). Recent evidence suggests that the CGRP receptor may also play a significant role in AD due to its involvement in neuroinflammatory pathways common to both conditions. Notably, Na et al., found that the CGRP receptor antagonist BIBN could alleviate AD-related pathologies in young 5×FAD mouse models (Na et al., 2020). Furthermore, several studies have identified a positive correlation between acute migraine attacks and an increased risk of dementia, based on long-term follow-up data from multiple countries, including Canada (Medrea et al., 2024), Korea (Hurh et al., 2022), and the United Kingdom (Kostev et al., 2019). This human-based association suggests a dual therapeutic potential of CGRP receptor inhibition in both migraine treatment and AD improvement.

In conclusion, this study demonstrates that inhibition of CGRP receptor mitigates AD pathology in 5×FAD mice, potentially by reprogramming neuronal lipids metabolism via the HDAC11/LXRβ/ABCA1 pathway, a mechanism conserved in humans. Our findings not only elucidate a novel mechanism in AD pathogenesis but also provide compelling evidence supporting the targeting of the CALCRL/HDAC11 axis as a promising therapeutic strategy for AD treatments.

## RESOURCE AVAILABILITY

All data reported in this paper will be shared by the lead contact upon request. This paper does not report original code.

## ACKNOWLEDGMENTS

This study was supported by National Natural Science Foundation of China (32271003), the Science and Technology Commission of Shanghai Municipality (24141901200), the Open Research Fund of Navy Medical University Basic Medical College (Grant number ORFBMC-JCKFKT-MS-015), the Shanghai Municipal Science and Technology Major Project (No. 2018SHZDZX01) and ZJLab, the Shanghai Center for Brain Science and Brain-Inspired Technology, the Innovative Research Team of High-Level Local University in Shanghai, STI2030-Major Project #2021ZD0201100 Task 4 #2021ZD0201104 from the Ministry of Science and Technology (MOST). We thank Meng Liu from the Core Facility of Basic Medical Sciences, Shanghai Jiao Tong University School of Medicine, for their invaluable contribution to the lipidomic analysis in this study.

## AUTHOR CONTRIBUTIONS

F.H., R.S., J.F., W.L., M.Y., and L.C. proposed and supervised the study. G.F. and F.C. performed most experiments and analyzed the data, with assistance from M.L., P.Y., H.D., J.S., P.N., C.S., H.L., J.Z., Y.Z., X.Y., X.T., M.W., Z.W., Z.C., H.L., X.L., R.Z., Y.L.. All authors contributed to the interpretation of data and the revision of the manuscript.

## DECLARATION OF GENERATIVE AI AND AI-ASSISTED TECHNOLOGIES

This study did not utilize generative AI or AI-assisted technologies in any part of the research process.

## STAR METHOD

### Mice

C57BL/6Smoc (wild type [WT]) mice and *Calca^-/-^* mice (C57BL/6Smoc-Calcaem1Smoc, Cat. NO.

NM-KO-190928) from Shanghai Model Organisms Center, Inc. (Shanghai, China), and 5×FAD mice (Tg(APPSwFlLon,PSEN1*M146L*L286V)6799Vas, MMRRC stock #34848-JAX)(Oakley H, et al. 2006) were used for experiments in this study and genotyping was performed using primers and polymerase chain reaction (PCR) conditions listed on the vendor website (Shanghai Model Organisms Center and JaxLab). All animals were maintained on the C57BL/6Smoc background, housed under a 12 h light/12 h dark cycle with libitum access to food and water. All experiments involving animal procedures were approved by the Institutional Animal Care and Use Committee of Fudan University, Shanghai Medical College.

### Caenorhabditis elegans

The C. elegans strains CL2006 (myo-3p::A-Beta (1-42)::let-851 3‘UTR)+ rol-6(su1006) and E. coli strain OP50 which were purchased from the Caenorhabditis Genetic Center (University of Minnesota, Minneapolis, MN). CL2006 were maintained at 20 °C, on solid nematode growth medium (NGM) plated with live OP50 as a food source.

### Human Subjects

The PART/ABC score distinguishes Primary Age-Related Tauopathy (PART) from Alzheimer’s disease by assessing A (amyloid-β deposition, Thal phase 0-5), B (neurofibrillary tangles, Braak stage 0-VI), and C (neuritic plaque density, CERAD score 0-3), where PART is characterized by tau pathology (B) in the absence of amyloid (A0).

For western blotting: hippocampal CA1 samples from adult humans with Alzheimer’s disease and age- and sex-matched controls were obtained from Fudan branch of National Health and Disease Human Brain Tissue Resource Center, the Body Donation Station in Fudan University (Shanghai Red Cross Society). Information on the samples are listed in Figure S1h. No differences were observed between genders. The use of anonymized excess tissue materials for scientific analyses was approved by the local ethics committee (Research Ethics Committee reference number: 2025- 003).

### Drug treatments

Solutions of Rimegepant (TargetMol Chemicals Inc) were prepared for intraperitoneally (i.p.) injection in mice, following a dosing regimen of 10 mg/kg every other day for a total of 15 doses.

sAβ oligomer was produced from β-Amyloid (1-42), (rat/mouse) (TFA) (TargetMol Chemicals Inc), which was administrated at a concentration of 10 μM for 24 h in primary cortical neurons from E14.5 embryos and mixglial culture from newborn C57BL/6Smoc or *Calca^-/-^* mice. Elevenostat (10 μM) (TargetMol Chemicals Inc) was administered via stereotactic injection (1 μL) in 5×FAD mice. All chemical powders were dissolved in Dimethyl Sulfoxide (DMSO, Beyotime Biotechnology) for storage.

### Viral vectors

rAAV-hSyn-EGFP-5’miR-30a-shRNA(Hdac11)-3’-miR30a-WPREs [5x10^12^ viral genome (v.g.)/mL] and rAAV-hSyn-EGFP-5’miR-30a-shRNA(scramble)-3’-miR30a-WPREs (5x10^12^ v.g./mL) were produced by BrainVTA(Wuhan) Co.,Ltd (Wuhan, China), and aliquoted and stored at 80 ℃ until use.

### Molecular docking

The optimized CGRP receptor 3D structure was acquired from PDB database (PDB ID: 3N7R). The pre-processing of the CGRP receptor 3D structure was implemented using Accelrys Discovery studio 2.5 (DS2.5). The docking procedure was implemented using the LibDock program of the DS2.5 simulation software package. The 3D structure of the Hdac11 (AF-Q91WA3-F1) and Nr1h2/LXRβ (AF-Q60644-F1-v4) obtained from AlphaFoldDB (https://alphafold.ebi.ac.uk/) was then minimized on DS2.5. The protein–protein docking simulations were carried out using the HDOCK server. PyMol 1.5 (Schrodinger LLC) was used for visual inspection of the results and graphical representations.

### Surgery and viral injection

Mice were anesthetized with isoflurane and restrained on a stereotaxic apparatus (RWD Life Science) by nose clip and ear bars. Erythromycin ointment (Guangdong Hengjian Pharmaceutical Co., Ltd) was used to moisten the corneal. The scalp was shaved and cleaned, and then a linear incision on the skin was made to expose the skull, which was subsequently cleaned with a cotton swab dipped in normal saline. The brain was adjusted to be flat in both x and y coordinates. According to “The Mouse Brain” (Keith B.J. Franklin and George Paxinos, the third edition, Elsevier), bregma was defined as the origin of stereotaxic with x = 0, y = 0, z = 0, and lambda was defined as x = 0, y = −4.2 mm, z = 0. If the distance between bregma and lambda was not 4.2 mm, the coordinates for the injection site were calculated proportionally. The skull above the target area was drilled open and a glass pipette (WPI, pulled by Sutter PC-10, NARISHIGE) was inserted into the injection site. 300 nL rAAV or 1 μL Elevenostat (10 μM) were bilaterally injected into CA1 regions of 5×FAD mice (AP, - 2.00 mm from bregma; ML, ± 1.60 mm; DV, - 1.90 mm from the skull) at the rate of 80 nL/min for each site. After injection, the glass pipette was pulled out slowly after 10-min staying.

### Paralysis time assay

Paralysis assay was performed in CL2006 worms to assess the effect of Rimegepant and Elevenostat on delaying the onset of amyloid-β-induced paralysis. Petri dishes (6cm) containing Nematode Growth Medium (NGM) were supplemented with Rimegepant at concentrations of 0.4 μM, 2 μM, or 10 μM, or Elevenostat at 4 μM, 20 μM, or 100 μM. After the agar had solidified, the compounds were allowed to diffuse throughout the agar and incubated at room temperature for 30 minutes. The Petri plates were then kept at 16 °C overnight and seeded with 50 μL OP50 (0.5 mg/mL) as previously described.

### Open field test (OF)

The animal spontaneous locomotor activity was measured in a plexiglass box (42×42×50cm), recorded by Ethovision XT14.0 software (Noldus). During testing, mice were placed in the center of the activity chamber and allowed to freely explore for 10 min, not affected by external stimuli. Anxiety-related performance was quantified by calculating the number of times entering the central area. The time spent in the center of the box (half of the box area) was also recorded.

### Novel location recognition test (NLR)and Novel object recognition test (NOR)

The novel location recognition test or novel object recognition test was performed in an open field where the movements of mice were recorded via a camera mounted above the field. Before each test, mice were acclimatized in the behavioral room for 30 min and were given a 10 min habituation trial with no objects on the open field.

First, each mouse was placed on one side of the open field and allowed to explore freely for 10 minutes. After one day, two identical objects were presented to the mouse for 10 min. The following day, one object was relocated to a novel position, and the mouse was allowed to explore for 10 min. On the subsequent day, one object was replaced with a novel object, and the mouse was given 10 min to explore. The chamber and objects were cleaned with 75% ethanol between trials to remove olfactory stimuli. Testing sessions were video monitored, and object exploration times were scored by a blinded experimenter. The results were quantified as the frequency of exploration for the novel object compared to the total frequency of novel and familiar object or location.

### Y-maze test

The Y-maze apparatus was made of acrylic plastic and was either black or gray in color. Mice were placed in the center of the Y-shaped maze, which consisted of three equal arms (A, B, C), each measuring 50 cm x 10 cm. Mice were allowed to explore the maze for 10 min, and the number and sequence of entries into the different arms were recorded. Three consecutive entries into different arms were considered one alternation. Percentage of alternation was calculated as follows: number of triads containing entries into 3 different arms/(total number of arms entered −2) × 100%. After each test, the mouse was carefully removed from the maze, and the maze was cleaned with 75% ethanol followed by distilled water. All other experimental conditions were kept consistent between the group.

### Elevated plus maze test (EPM)

The elevated plus maze consists of two open arms (without walls, 35 cm long X 6 cm wide) and two closed arms (with walls 19 cm tall), the intersection of the arms is 6 cm x 6 cm wide, and the entire maze is elevated 73 cm above the ground. Mice were first allowed to habituate in the testing room under dim light for 1 h prior to the start of testing. During testing, mice are placed in the maze at the intersection of the open and closed arms and allowed to freely explore the maze for 10 min. The maze is cleaned with 75% alcohol between testing of each mouse. The anxiety index was measured as follows: 1 − [(time spent in open arm (OA)/total time on the maze) + (number of entries to the OA/total exploration on the maze)/2] (Cohen et al., 2012).

### Forced swimming test (FST)

Forced swimming test was also for assessing depressive-like behavior. Each mouse was placed individually into glass cylinders. The test lasted for 6 min and the activity of mouse was video-recorded for analysis. Immobility was defined as the absence of any movement except for the head above water.

### Morris water maze test (MWM)

Mice were trained to search an unhidden circular escape platform (10 cm) in a pool (120 cm diameter) filled with opaque water by using different cues. The pool was filled with tap water mixed with non-toxic white paint. During the whole test the water temperature was maintained at 20 ± 2 °C. The pool was divided into four virtual quadrants based on the platform localization: left, right, opposite, and target.

Each mouse went through 5 or 6 days of acquisition training and a final probe trial. During the acquisition training, the platform was not hidden under the water. Mice were introduced into the water near the edge of the pool facing the wall. They were given one minute to find the submerged platform. The platform was located in the upper third of the quadrant (10 cm from the rim of the pool). If they failed to find the platform in one minute, they were gently guided to it. Every mouse had to sit on the platform for 15 s before being removed from the pool. To prevent hypothermia mice were kept in front of a heat lamp until they were dry. The cued training consisted of four trials per day with an average inter-trial interval of 60 min. The start position changed for every trial with the stationary location of platform. Twenty-four hours after the last acquisition trial, a probe trial was performed to assess spatial reference memory. For the probe trial the platform was removed from the pool, and mice were introduced into the water from a novel entry point (opposite of the platform location). Mice were allowed to swim freely for one minute. To calculate the time to reach the target, the position of the platform in the tracking software was set to the same position as in the previous acquisition training. For tracking the learning curve in training phases, platform crossing number, and percentage time spent in target quadrant, Ethovision XT14 video tracking software (Noldus) was used.

### Immunohistochemistry and immunofluorescence microscopy

Brain tissue/slices were post-fixed in 4% PFA in PBS at 4 °C overnight then subsequently transferred into 30% sucrose solution until cryo-sectioning. After embedding in optimal cutting temperature (OCT) compound, the slices were further cut coronally or sagittally into 30-μm-thick sections on a cryostat (Leica Biosystems). Immunohistochemistry and immunofluorescence protocols used were as previously published (Dong et al., 2024). For immunofluorescence (IF) staining, 30 μm thick free-floating brain sections or sections mounted on glass slides were washed, blocked and permeabilized by incubating in TBS containing 0.30% Triton X-100 and 5% horse serum for 1h at room temperature. Primary antibodies were diluted in TBS containing 0.30% Triton-X-100 and 1% horse serum. After overnight incubation at 4 °C with primary antibodies, the sections were rinsed 3x in TBS containing 1% horse serum at room temperature for 10 min each. Then, the sections were incubated in the appropriate fluorophore-conjugated secondary antibody at room temperature for 2 h in the dark. The sections were rinsed once and incubated with DAPI (1µg/ml) for 5min, washed 3x in TBS for 10 min, dried, and cover slipped with ProLong Diamond Antifade Mountant (Thermo Fisher).

### Thioflavin-S, Fluoro-Jade B and BODPIY staining

For Aβ plaques staining, briefly deparaffinized and rehydrated slides were incubated for 10 min in 1% thioflavin-S (Sigma) which was dissolved in 50% ethanol and the slides were subsequently rinsed in 80% ethanol and 50% ethanol. For Fluoro-Jade B staining detecting neuronal death, following immunofluorescence staining and subsequent drying for 3 h at room temperature, the brain sections were rehydrated for 2min and then stained with a solution of 0.06% potassium permanganate for an additional 2 min. Following this, the sections were washed and subsequently stained with 0.0001 % FJB solution, which contained 0.1% acetic acid, for a duration of 10min. For detecting lipid droplets, BODIPY™ 493/503 (Thermo Fisher) was dissolved in PBS to prepare a 10 μM working solution. Sections were imaged using a Nikon AIR-MP confocal microscope (Japan). Image-J was used to quantify and analyze the number of plaques, FJB+ neurons, BODIPY signals, and the percent area of BODIPY signals in cells.

### Preparation of soluble amyloid β 1-42 Oligomers (sAβ1-42)

First, lyophilized synthetic murine Aβ1-42 was dissolved in 100% HFIP (hexafluoroisopropanol) (Sigma-Aldrich) and incubated at room temperature for 1 h to establish randomization of structure formation. The HFIP was then evaporated, and the dry peptide was stored at -20 °C. For oligomeric conditions, the peptide was resuspended in DMSO to a final concentration of 10mM, then diluted in cell culture medium to achieve the required concentration of 1 mM and incubated at 4°C for 24 h before use.

### Scanning electron microscopy

The samples for scanning electron microscopy (SEM) analysis were prepared using the following method: 100 μL of sAβ1-42(1 mM) was centrifuged at 13,000g for 5 min and then applied to the polished surface of a silicon wafer and air-dried. The samples were analyzed using a desktop field-emission SEM (Phenom Pharos-SED-EDS G2, Thermo Fisher) at an acceleration voltage of 10 kV.

### Primary mixglia and cortical neuronal culture

For primary mixglial culture, the brains of WT mice at P1-P2 were dissected, and the meninges were removed in Hanks’ balanced salt solution (HBSS). The brains were then mechanically dissociated into single-cell suspension within HBSS. After centrifugation, the cells were incubated in Dulbecco ’s modified eagle medium (DMEM) containing 10% fetal bovine serum (FBS) in a 25 cm^2^ flask with 5% CO2 at 37 ◦C. The culture media were changed 24 hours later and then twice a week. The AD-like treatment and intervention or medication could be proceeded after 21-day culture.

For primary cortical neuronal culture, the pregnant female mice at E15.5-17 were terminally anesthetized with isoflurane. Abdominal skin was washed with 75% ethanol followed by an incision to cut and open the peritoneal cavity. Amniotic sacs were exposed with fine scissors and embryos were taken out from the uterus. The brains of embryos were dissected, and the meninges were removed in HBSS. The tissues were then thoroughly snipped and trypsinized (0.05% trypsin) at 37◦C for 8-10 min to obtain a single cell suspension. The suspension was centrifuged, and the cells were incubated in DMEM containing 10% FBS in a 24-well plate under 5% CO2 at 37 ◦C. The culture media were changed to neurobasal containing 2% B27 and 1% GlutaMax 4h later and then half-changed every 72 h. After 7 days, the cortical neurons would be ready good enough for experimental use.

### Cell viability assays

Cell viability was assessed using the Cell Counting Kit-8 (CCK-8) (Beyotime) and Hoechst 33342/PI double staining kit (Beyotime). Briefly, cells were seeded in 96-well or 24-well plates and allowed to grow to 80% confluence. Cells were then treated with sAβ1-42 (10 µM) with or without various doses of Rim or Ele for 24 h, with sAβ1-42 vehicle as the control. Cell viability was determined using CCK-8 and Hoechst 33342/PI staining according to the manufacturer’s instructions.

### Western blotting

Total protein was extracted from cultured cells and tissues using RIPA Lysis Buffer (Beyotime) supplemented with protease inhibitor cocktail (Beyotime). The protein samples were then separated by SDS-PAGE and transferred onto PVDF membranes (Millipore). For HRP-conjugated secondary antibodies, proteins were detected using the enhanced chemiluminescence method (ECL) (BioRad). The quantification of each band was performed by densitometric measurement using ImageJ software.

### Immunoprecipitation and Immunoblotting

Immunoprecipitation (IP) assays were performed using the Pierce™ Classic Magnetic IP/Co-IP Kit (Thermo Fisher), following the manufacturer’s protocol. Briefly, hippocampal tissue was lysed in RIPA Lysis Buffer (Beyotime) for 20 min on ice. The cell lysates were then centrifuged at 13,000 rpm for 15 min, and the supernatants were incubated overnight with Anti-HDAC11 antibody or Anti-acetyl-Lysine antibody. After overnight incubation, protein A/G magnetic beads (washed three times with IP Lysis/Wash Buffer) were added to the cell extracts and incubated with gentle shaking for 1 hour at room temperature. The beads were then washed three times with IP Lysis/Wash Buffer, resuspended in 50 μL of SDS sample buffer, and boiled at 97 °C for 10 min. The immunoprecipitates were subsequently analyzed by standard immunoblotting.

### Elisa for CALCA

Primary cortical neurons were prepared as described in the Primary Microglia and Cortical Neuron Culture section. Briefly, cells were seeded in 24-well plates and allowed to grow to 80% confluence. Cells were then treated with sAβ1-42 (10 µM), with the sAβ1-42 vehicle used as the control. After determining total protein concentrations using the BCA assay, the concentration of CALCA in both the supernatant and cell lysates was measured using an ELISA kit for CALCA (Cloud-Clone Corp), following the manufacturer’s protocol.

### Data analysis of snRNA-seq datasets

snRNA-seq dataset (GSE148822) contains 18 samples including 482,472 sequenced nuclei with 10 AD samples including 3 male and 7 female AD donors, 5 control without amyloid plaques (CTR) samples and 3 control plus amyloid plaques (CTR+) samples. The expression matrix was imported into R (version 4.3.1) using the Seurat package (version 2.3.4). Default parameters within the Seurat package were employed unless otherwise specified. Cells were excluded if they exhibited fewer than 200 or more than 5000 genes, or if more than 10% of their expressed genes were mitochondrial, to ensure that only cells in good condition and without doublets were included. After independent normalization of the data, we merged the patient and control expression matrixes using Seurat built-in function called “Run Harmony”. This was followed by scaling the matrix and performing principal component analysis (PCA) based on the top 2000 highly variable genes. The top 11 principles were employed to construct uniform manifold approximation and projection (UMAP).

### RNA extraction, RT-qPCR and RNA-seq

Total RNA was extracted from cultured cells and the tissues using TRIzol reagent (TIANGEN), following the manufacturer’s protocol. Reverse transcription was carried out using random primers and the Fastking gDNA Dispelling RT SuperMix (TIANGEN). Quantitative real-time PCR using the SuperReal PreMix Plus (SYBR Green) (TIANGEN).

For RNA-seq, the quality of the total RNA was first assessed. Then, the library was constructed, purified, detected and quantified. Following the generation of sequencing clusters, sequencing was conducted on the Illumina/MGI platform.

### RNA-seq

After quality control, the raw data were aligned to the mice reference genome. Differential expression analysis was performed using the DESeq (2012) R package. adj. *p* value <= 0.05 was set as the threshold for significantly differential expression. GOseq was employed to analyze the GO enrichment of the differentially expressed genes in molecular function, cellular component, and biological process, and the criteria for screening significant enrichment were over represented adj. *p* value <= 0.05. GSEA was performed using the “GSEA” R package. analysis was WGCNA analysis was realized with R package “WGCNA” (version 1.68). The clustering of the genes was performed with “Mfuzz” R package.

### Protein digestion and desalination

Using QLBIO MAGicomicomicS-MMB8X, 20 μL of protein was added to eight tandem tubes containing MMB beads and incubated at 37 °C for 30 min. After incubation, 45 μL of bonding solution was added and incubated at room temperature for 15 min. Following this, the supernatant was discarded, and the MMB beads were washed three times with cleaning solution. The magnetic beads were then re-suspended in 40 μL of enzyme working solution and incubated at 37 °C for over 4 h. After incubation, 5 μL of termination solution was added to halt the enzyme reaction, and the mixture was freeze-dried.

### LC-MS/MS analysis

Nanoflow LC-MS/MS analysis of tryptic peptides was performed on a quadrupole Orbitrap mass spectrometer (Thermo Fisher) coupled to an EASY nLC 1200 ultra-high pressure system (Thermo Fisher) via a nano-electrospray ion source. One microgram of peptides was loaded onto a 25 cm column (100 μm inner diameter, packed with ReproSil-Pur C18-AQ 1.5-µm silica beads; QL- HPLC-100*15; Beijing Qinglian Biotech Co., Ltd). Peptides were separated using a gradient: 8–12% B in 2min, followed by 12–30% B in 3min, and then increased to 40% B in 7min, followed by a 10min wash at 95% B at a flow rate of 300nL per minute. Solvent A was 0.1% formic acid in water, and solvent B was 80% acetonitrile (ACN) with 0.1% formic acid in water. The total run duration was 30 min. The column temperature was maintained at 60 °C using an in-house-developed oven. The mass spectrometer was operated in “top-40” data-dependent mode, collecting MS spectra in the Orbitrap mass analyzer (120,000 resolution, 350–1500 m/z range) with an automatic gain control (AGC) target of 3E6 and a maximum ion injection time of 80 ms. The most intense ions from the full scan were isolated with an isolation width of 1.6m/z. Following higher-energy collisional dissociation (HCD) with a normalized collision energy (NCE) of 27, MS/MS spectra were acquired in the Orbitrap (15,000 resolution) with an AGC target of 5E4 and a maximum ion injection time of 45 ms. Precursor dynamic exclusion was enabled with a duration of 16s.

### The identification and quantitation of protein

All raw files were analyzed using the Proteome Discoverer suite (Thermo Fisher). MS2 spectra were searched against the UniProtKB human proteome database, including Swiss-Prot human reference protein sequences. The Sequest HT search engine was used, with the following parameters: fully tryptic specificity, a maximum of two missed cleavages, a minimum peptide length of 6, fixed carbamidomethylation of cysteine residues (+57.02146 Da), variable oxidation of methionine residues (+15.99492 Da), a precursor mass tolerance of 15 ppm, and a fragment mass tolerance of 0.02 Da for MS2 spectra collected in the Orbitrap. A Percolator was used to filter peptide spectral matches and peptides to a false discovery rate (FDR) of less than 1%. After spectral assignment, peptides were assembled into proteins, which were further filtered based on the combined probabilities of their constituent peptides, maintaining a final FDR of 1%. By default, the top matching protein, or ‘master protein,’ is the protein with the largest number of unique peptides and the smallest value in percent peptide coverage (i.e., the longest protein). Only unique and razor (i.e., parsimonious) peptides were considered for quantification.

### Tissue lipid extraction

Forty microliters of cold water were added to accurately weigh 10 ± 0.2 mg of hippocampal tissue, followed by homogenization at 20 Hz for 2 min. Next, 20 µL of the supernatant was removed, and 225 µL of MeOH was added, followed by vortexing for 10 s. Then, 5 µL of the internal standard and 750 µL of MTBE were added, and the mixture was vortexed for 10 seconds both before and after adding MTBE. The mixture was then incubated for 30 min, followed by the addition of 188 µL of H₂O and vortexing for 10 s to form a two-phase system. After equilibration for 10 min at 4 °C, the mixture was centrifuged at 10,000g for 10 min at 4 °C. Seven hundred microliters of the supernatant were dried under nitrogen. All samples were reconstituted in ACN/IPA/H₂O (65:30:5, v/v/v), and 20 µL of the mixture was used as a quality control (QC) sample. Finally, 2 µL of each sample was injected into the LC-MS system.

### Proteomics analysis

Total protein from mouse hippocampal tissues was extracted. Proteins were labeled with iTRAQ labeling reagents (ABSCIEX, 4,381,663) and then subjected to Liquid Chromatography and tandem MS analysis. Protein analysis and relative iTRAQ quantification services were undertaken by OE Biotech. Differentially expressed protein (DEP) was performed using the DESeq R package. adj. *p* value <= 0.05 was set as the threshold for significantly differential expression. “KEGG” R package was used for KEGG pathway enrichment analyses of DEPs. The clustering of the proteins was performed with “Mfuzz” R package.

### Lipidomics

Lipidomic profiling was performed using an Ultra-Performance Liquid Chromatography-Mass Spectrometry (UPLC-MS/MS) system (UPLC, Agilent 1290; MS, Applied Biosystems SCIEX 6500+QTRAP) at the Core Facility of Basic Medical Sciences, Shanghai Jiao Tong University School of Medicine (Shanghai, China). The samples were separated using reverse-phase chromatography (Kinetex® 2.6 μm C18, 2.1 × 100 mm, Phenomenex). The experiment was conducted in Multi-Reaction Monitoring (MRM) mode, with each lipid type calibrated using corresponding lipid isotope standards to ensure experimental accuracy. Mass spectrometric data were processed using Analyst 1.6.3 software (AB Sciex). Data processing involved Sciex OS software, and the analytical data consisted of retention time (RT), normalization by internal standards (ISTDs), and baselining towards blank runs to remove background noise. After PCA, differential expression analysis was performed using the DESeq (2012) R package. adj. *p* value <= 0.05 was set as the threshold for significantly differential expression.

### Statistics and reproducibility

All data were analyzed using R studio (4.3.1) and differences between groups were tested with unpaired t-test, one-way or two-way analysis of variance (ANOVA) followed by Bonferroni multiple comparisons as indicated. Data are presented as the mean ± SEM. Significance levels were defined as follows: \*\*\*\**p* < 0.0001, \*\*\**p* < 0.001, \*\**p* < 0.01, \**p* < 0.05. All samples or animals were included in the statistical analysis, the value of n per group is included under figure legends.

