## Supplemental figure for "Inhibition of CGRP receptor ameliorates AD pathology by reprogramming lipid metabolism through HDAC11/LXRβ/ABCA1 signaling"

### SUPPLEMENTAL INFORMATION TITLES AND LEGENDS

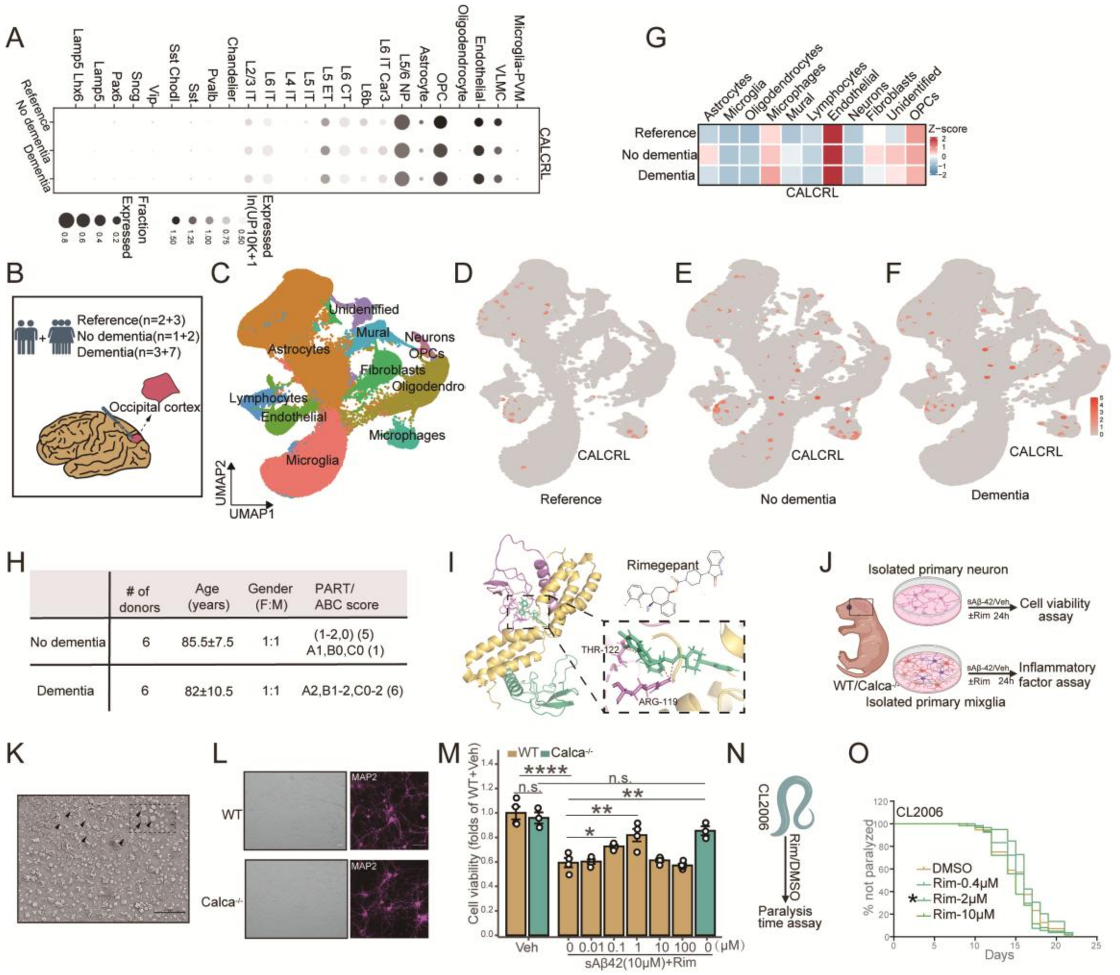


**Figure S1. CALCRL distribution in the brain and its inhibition alleviates sAβ1-42-induced neuronal cell death *in vitro* and reduces paralysis in *Caenorhabditis elegans* CL2006.**

(A) CALCRL expression dot plot within each sample.

(B) Characteristic information and the number of subjects in each group.

(C) UMAP of the associated cell type and clusters.

(D-F) UMAP visualization of cells expressing CALCRL in the reference (D), no dementia (E), and dementia (F) groups.

(G) CALCRL expression heatmap within each sample.

(H) Detailed characteristic information of subjects in each group.

(I) Chemical structure of Rim and its docking with the crystallized structure of CGRP receptor with PDB code 3N7R.

(J) Schematic representation of the experimental design.

(K) Representative scanning electron microscopy images of sAβ1-42.

(L) Cell identification of primary cortical neuron was determined with bright field and MAP2 immunofluorescence staining.

(M) The effect of different concentrations of Rim and *Calca* deficiency on primary cortical and hippocampal neuronal cell viability treated with sAβ1-42 estimated by CCK-8 assay.

(N) Schematic representation of the experimental design.

(O) Aβ-induced paralysis was delayed in AD transgenic nematode CL2006 treated with different concentrations of Rim. The numbers of worms for assays were 54–61 in each treatment. n=3-4/treatment. Scale bars: 40 μm and 1 μm (insert) in (K); 100 μm (left) and 50 μm (right) in (L). **p* < 0.05, ***p* < 0.01, *****p* < 0.0001. n.s., non-significant. Statistical analyses were analyzed by one-way ANOVA followed by a post hoc Dunnett's test (M), and Gehan-Breslow-Wilcoxon Test (O).

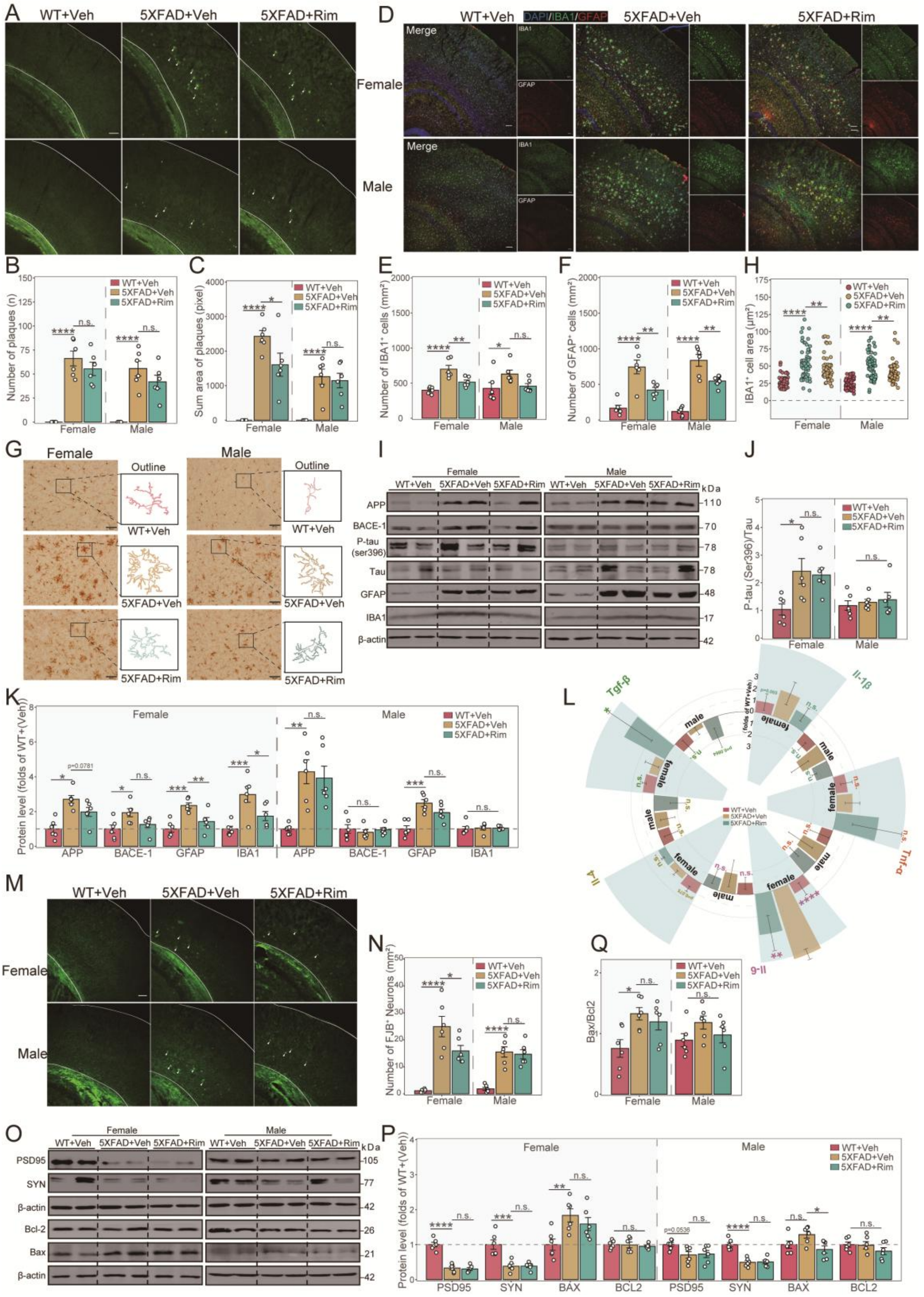


**Figure S2. Rim alleviates AD-related pathological features in the cortex.**

(A) Representative Thioflavin-S staining cortical images of WT and 5×FAD mice after the administration of the vehicle or Rim.

(B and C) Quantitative measurements of Aβ plaque numbers (B) and area (C) in the cortex. (D) Immunofluorescence staining of GFAP (red) and IBA-1 (green) in the cortex.

(E and F) Quantifications of $GFAP^+$ astrocytes (E) and IBA-$1^+$ microglia (F) in the cortex.

(G and H) Representative images of IBA-1 immunostaining (G) and the quantified IBA-$1^+$ cell body volume (H) in the cortex.

(I-K) Western blot image (I) and densitometry analysis of AD-related marker proteins including APP, BACE1, phosphorylated tau, total tau, GFAP and IBA-1 in the cortex (J and K).

(L) qPCR analysis of pro-inflammatory factors Il-1β, Tnf-α, and Il-6, and anti-inflammatory factors Il-4 and Tgf-β expression in the cortex.

(M) Representative Fluoro-Jade B staining images.

(N) Quantitative measurements of $FJB^+$ neuronal numbers.

(O-Q) Western blot image (O) and densitometry analysis of PSD95, synaptophysin (SYN), Bcl-2 and Bax in the cortex (P and Q). n=3-6/genotype; n=60 IBA-$1^+$ cell, 6/group (H). Scale bar: 100 μm in (A, B and M); 200 μm in (G). $*p < 0.05$, $**p < 0.01$, $***p < 0.001$, $****p < 0.0001$, n.s., non-significant. Statistical analyses were analyzed by one-way ANOVA followed by a post hoc Dunnett's test.

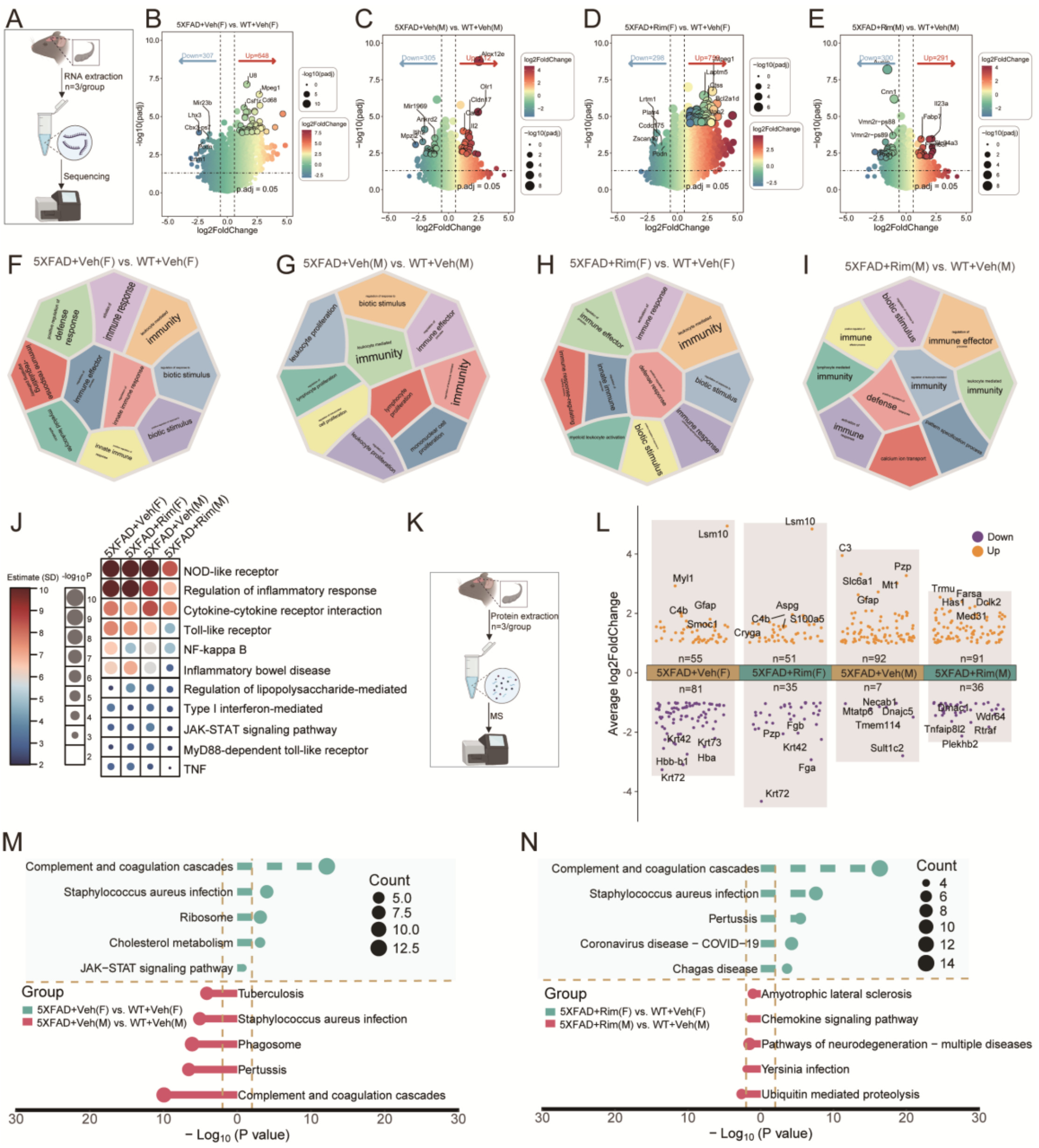


**Figure S3. Functional enrichment analysis of differentially expressed genes and proteins.**

(A) Schematic representation of the experimental design.

(B) Volcano plot of differentially expressed genes (DEGs) in 5×FAD+Veh(F) vs. WT+Veh(F) mice.

(C) Volcano plot of DEGs in 5×FAD+Veh(M) vs. WT+Veh(M) mice.

(D) Volcano plot of DEGs in 5×FAD+Rim(F) vs. WT+Veh(F) mice.

(E) Volcano plot of DEGs in 5×FAD+Rim(M) vs. WT+Veh(M) mice.

(F) Gene Ontology (GO) enrichment analysis of the top 10 biological process of the DEGs in 5×FAD+Veh(F) vs. WT+Veh(F) mice.

(G) GO enrichment analysis of the top 10 biological process of the DEGs in 5×FAD+Veh(M) vs. WT+Veh(M) mice.

(H) GO enrichment analysis of the top 10 biological process of the DEGs in 5×FAD+Rim(F) vs. WT+Veh(F) mice.

(I) GO enrichment analysis of the top 10 biological process of the DEGs in 5×FAD+Rim(M) vs. WT+Veh(M) mice.

(J) Gene set enrichment analysis (GSEA) of inflammatory pathways of DEGs in each group, vs. Corresponding WT+Veh group.

(K) Schematic representation of the experimental design.

(L)Multiple Volcano plots of differentially expressed proteins (DEPs) in each group, vs. Corresponding WT+Veh group.

(M) Kyoto encyclopedia of genes and genomes (KEGG) pathways enrichment analysis of DEPs in 5×FAD+Veh(F) vs. WT+Veh(F) and 5×FAD+Veh(M) vs. WT+Veh(M) mice.

(N) KEGG pathways enrichment analysis of DEPs in 5×FAD+Rim(F) vs. WT+Rim(F) and 5×FAD+Rim(M) vs. WT+Rim(M) mice.

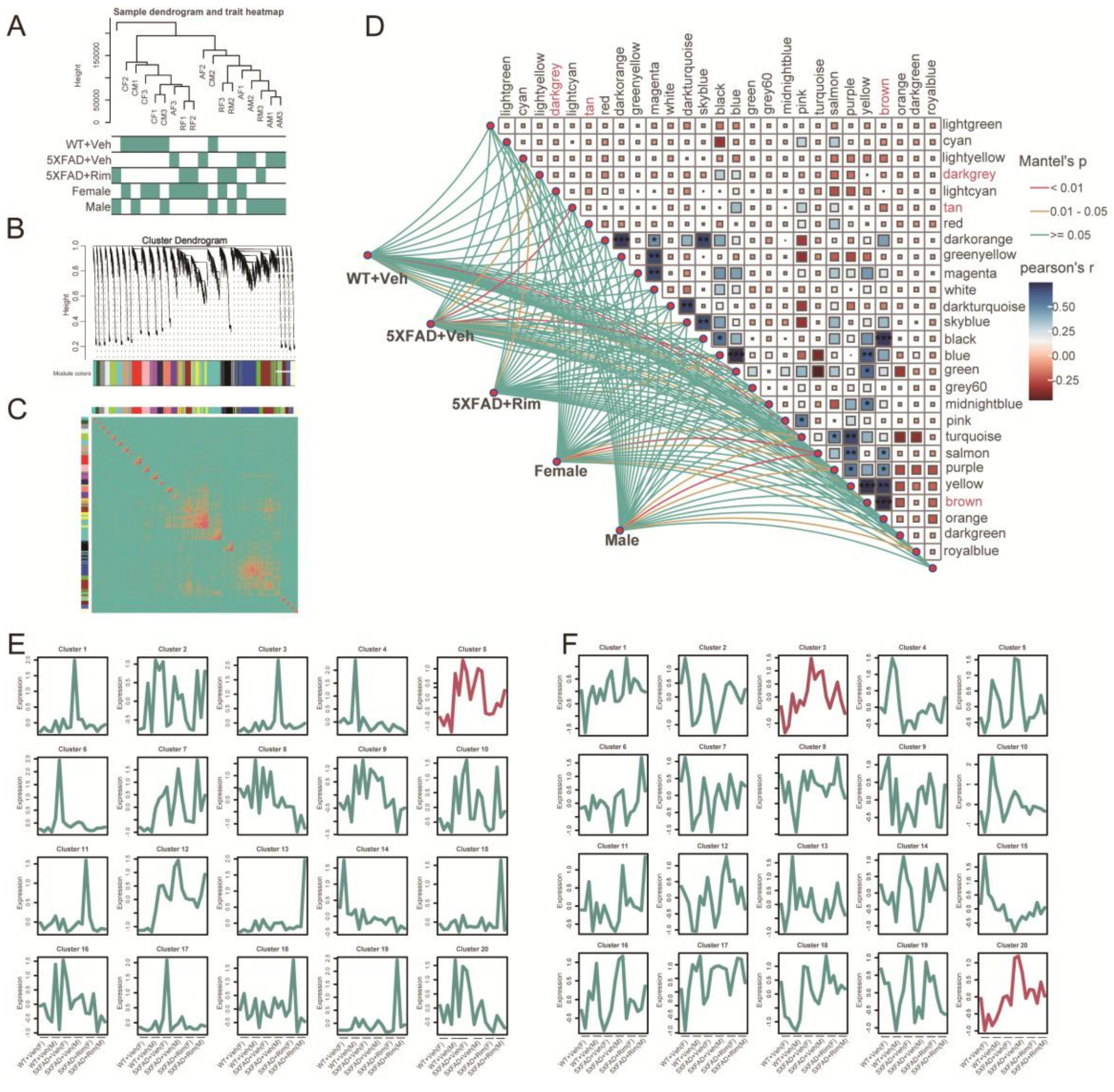


**Figure S4. Weighted gene co-expression network analysis (WGCNA) and gene expression trend analysis.**

(A) Sample dendrogram.

(B) Module gene clustering tree graph.

(C) Module gene correlation heat map.

(D) Heatmap for the relationships of modules and trait.

(E) Soft clustering of the expression of genes using the Mfuzz algorithm.

(F) Soft clustering of the expression of proteins using the Mfuzz algorithm.

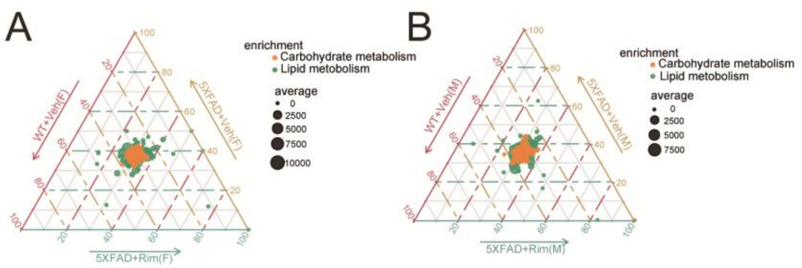


**Figure S5. Ternary plot analysis of gene expression related to lipid and carbohydrate metabolism.**

(A and B) Ternary plot analysis of gene expression related to lipid and carbohydrate metabolism in female (A) and male (B) 5×FAD mice.

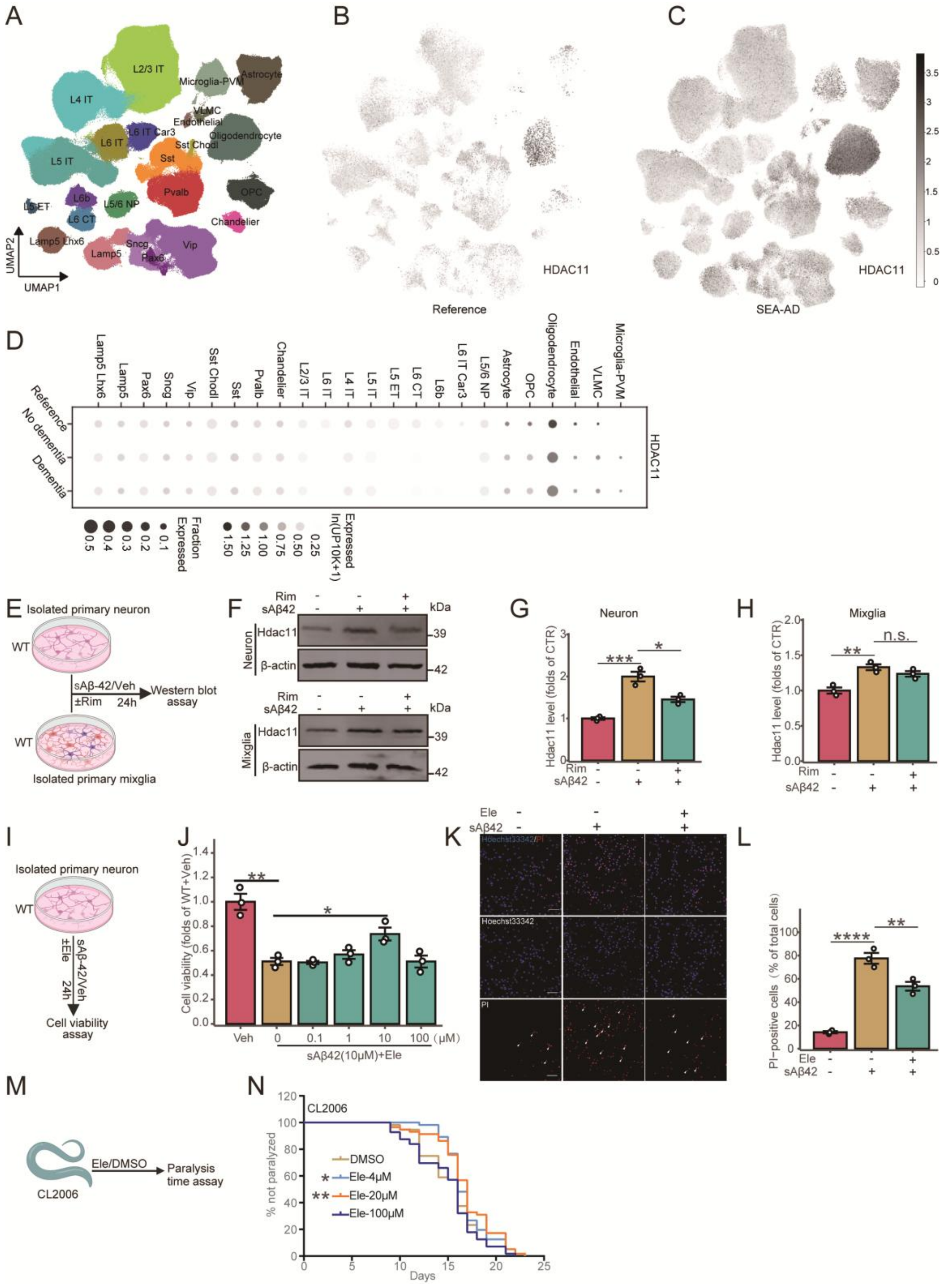


**Figure S6. HDAC11 distribution in the brain and its inhibition alleviates sAβ1-42-induced neuronal cell death *in vitro* and reduces paralysis in *Caenorhabditis elegans* CL2006**

(A) UMAP of the associated cell type and clusters.

(B and C) UMAP visualization of cells expressing HDAC11 in the reference (B) and SEA-AD (C) groups.

(D) HDAC11 expression dot plot within each sample.

(E) Schematic representation of the experimental design.

(F-H) Western blot image (F) and densitometry analysis of HDAC11 in primary neuron (G) and mixglia (H).

(I)Schematic representation of the experimental design.

(J) The effect of different concentrations of Ele on primary cortical neuron cell viability treated with sAβ1-42 estimated by CCK-8 assay.

(K and L) Representative images of Hoechst 33342 and PI double fluorescent staining (K) and the quantification of PI-positive cells (L).

(M) Schematic representation of the experimental design.

(N) Aβ-induced paralysis was delayed in AD transgenic nematode CL2006 treated with different concentrations of Ele. The numbers of worms for assays were 54–61 in each treatment. n=3/treatment. Scale bars: 50 μm in (K). *$p < 0.05$, **$p < 0.01$, ***$p < 0.001$, ****$p < 0.0001$. n.s., non-significant. Statistical analyses were analyzed by one-way ANOVA followed by a post hoc Dunnett's test (G-L), and Gehan-Breslow-Wilcoxon Test (N).

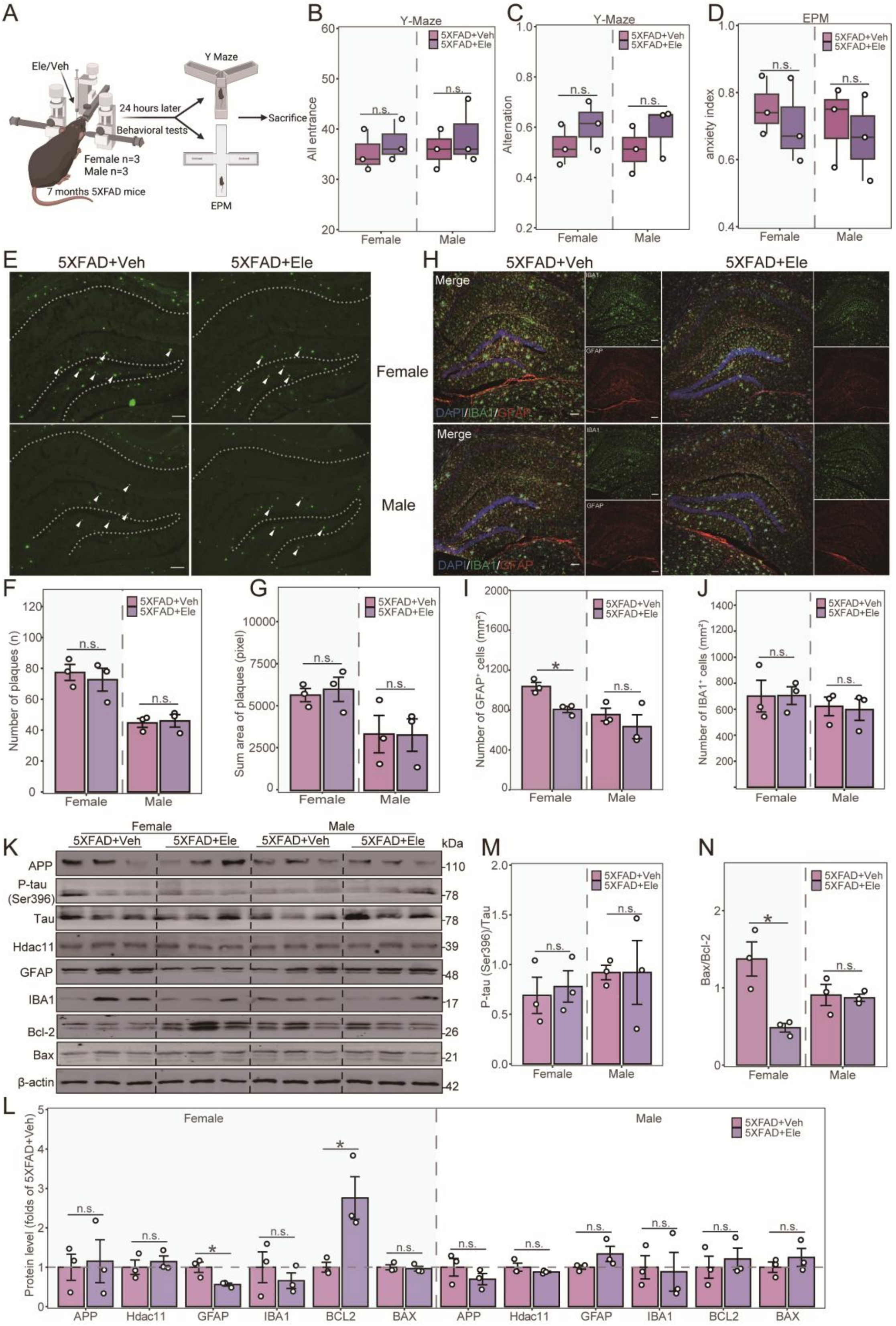


**Figure S7. Short-term inhibition of hippocampal HDAC11 function reduces astroglial activation in female 5×FAD mice.**

(A) Schematic representation of the experimental design.

(B and C) The number of total arm entries (B) and spontaneous alternations (C) in the Y-maze test.

(D) The anxiety index calculated by the EPM.

(E) Representative Thioflavin-S staining hippocampal images of 5×FAD mice after the administration of the vehicle or Ele.

(F and G) Quantitative measurements of Aβ plaque numbers (F) and area (G) in hippocampus. (H) Immunofluorescence staining of GFAP (red) and IBA-1 (green) in the hippocampus.

(I and J) Quantifications of $GFAP^+$ astrocytes (I) and $IBA\text{-}1^+$ microglia (J) in hippocampus.

(K-N) Western blot image (K) and densitometry analysis of APP, phosphorylated tau, total tau, GFAP, IBA-1, Bcl-2 and Bax in the hippocampus (L-N). n=3/genotype. Scale bars: 100 μm in (E and H). $*p < 0.05$, n.s., non-significant. Statistical analyses were analyzed by unpaired Students' t-test.

A

| Rank | 1 | 2 | 3 | 4 | 5 | 6 | 7 | 8 | 9 | 10 |
|---|---|---|---|---|---|---|---|---|---|---|
| Confidence score | 0.9569 | 0.9562 | 0.8949 | 0.8922 | 0.8852 | 0.8796 | 0.8769 | 0.8767 | 0.8707 | 0.8634 |
| Docking score | -304.95 | -304.17 | -257.08 | -255.66 | -252.14 | -249.42 | -248.16 | -248.07 | -245.37 | -242.19 |

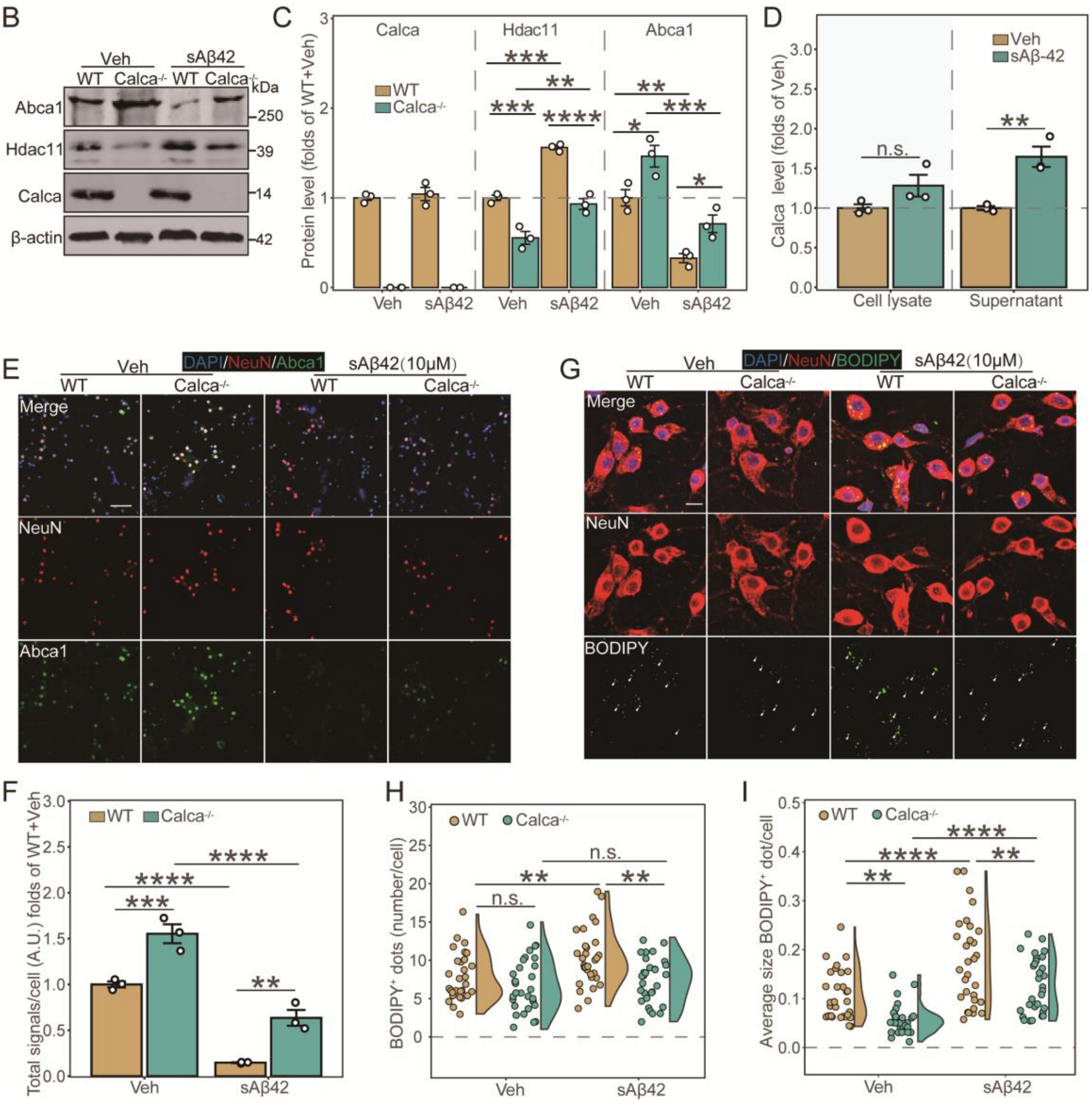


**Figure S8. Knockdown of *Hdac11* exerts a protective role *in vitro* by targeting LXRβ/ABCA1**

**axis.**

(A) The top 10 confidence score and docking score.

(B and C) Western blot image (B) and densitometry analysis of Abca1, Hdac11 and Calca in primary neuron isolated from WT and *Calca*$^{-/-}$ mice (C).

(D) ELISA analysis of α CGRP both in the cell lysates and the supernatants following sAβ1-42 treatment.

(E and F) Immunofluorescence staining (E) of NeuN (red) and Abca1 (green), and quantifications (F) in primary neurons isolated from WT and *Calca*$^{-/-}$ mice treated with or without sAβ1-42 for 24 hours.

(G-I) Immunofluorescence staining (G) of NeuN (red) and BODIPY (green), and quantifications dots numbers (H) and area (I) in primary neurons isolated from WT and *Calca*$^{-/-}$ mice treated with or without sAβ1-42 for 24 hours. n=3/genotype/treatment (C, D and F); n=30 NeuN$^{+}$ cell, 3 treatment/group (H and I). Scale bars: 100 μm in (E), 10 μm in (G). *$p < 0.05$, **$p < 0.01$, ****$p < 0.0001$. Statistical analyses were analyzed by Two-way ANOVA followed by Bonferroni post hoc analysis.
